# Cohort-wide deep whole genome sequencing and the allelic architecture of complex traits

**DOI:** 10.1101/283481

**Authors:** Arthur Gilly, Daniel Suveges, Karoline Kuchenbaecker, Martin Pollard, Lorraine Southam, Konstantinos Hatzikotoulas, Aliki-Eleni Farmaki, Thea Bjornland, Ryan Waples, Emil V. R. Appel, Elisabetta Casalone, Giorgio Melloni, Britt Kilian, Nigel W. Rayner, Ioanna Ntalla, Kousik Kundu, Klaudia Walter, John Danesh, Adam Butterworth, Inês Barroso, Emmanouil Tsafantakis, George Dedoussis, Ida Moltke, Eleftheria Zeggini

**Author notes:** These authors contributed equally (Arthur Gilly, Daniel Suveges, Karoline Kuchenbaecker).

## Abstract

The role of rare variants in complex traits remains uncharted. Here, we conduct deep whole genome sequencing of 1,457 individuals from an isolated population, and test for rare variant burdens across six cardiometabolic traits. We identify a role for rare regulatory variation, which has hitherto been missed. We find evidence of rare variant burdens overlapping with, and mostly independent of established common variant signals (*ADIPOQ* and adiponectin, *P*=4.2×10^−8^; *APOC3* and triglyceride levels, *P*=1.58×10^−26^; *GGT1* and gamma-glutamyltransferase, *P*=2.3×10^−6^; *UGT1A9* and bilirubin, *P*=1.9×10^−8^), and identify replicating evidence for a burden associated with triglyceride levels in *FAM189A* (*P*=2.26×10^−8^), indicating a role for this gene in lipid metabolism.

---

The role of rare sequence variants in the genetic architecture of medically-relevant complex traits is not well-understood. Population-scale deep whole genome sequencing can capture genetic variation across the entire allele frequency spectrum traversing the coding and non-coding genome. Here, to improve our understanding of the role of rare variants, we perform cohort-wide deep whole genome sequencing of 1,457 individuals from a deeply-phenotyped, isolated population from Crete, Greece (the HELIC-MANOLIS cohort^1–3^) at an average depth of 22.5x (Supplementary Figure 1), capturing 98% of true single nucleotide variants (SNVs) (Online Methods and Supplementary Figure 2). We address open questions on whole genome sequencing study design, analysis and interpretation, and identify burdens of coding and regulatory rare variants associated with cardiometabolic traits.

**Figure 1.**
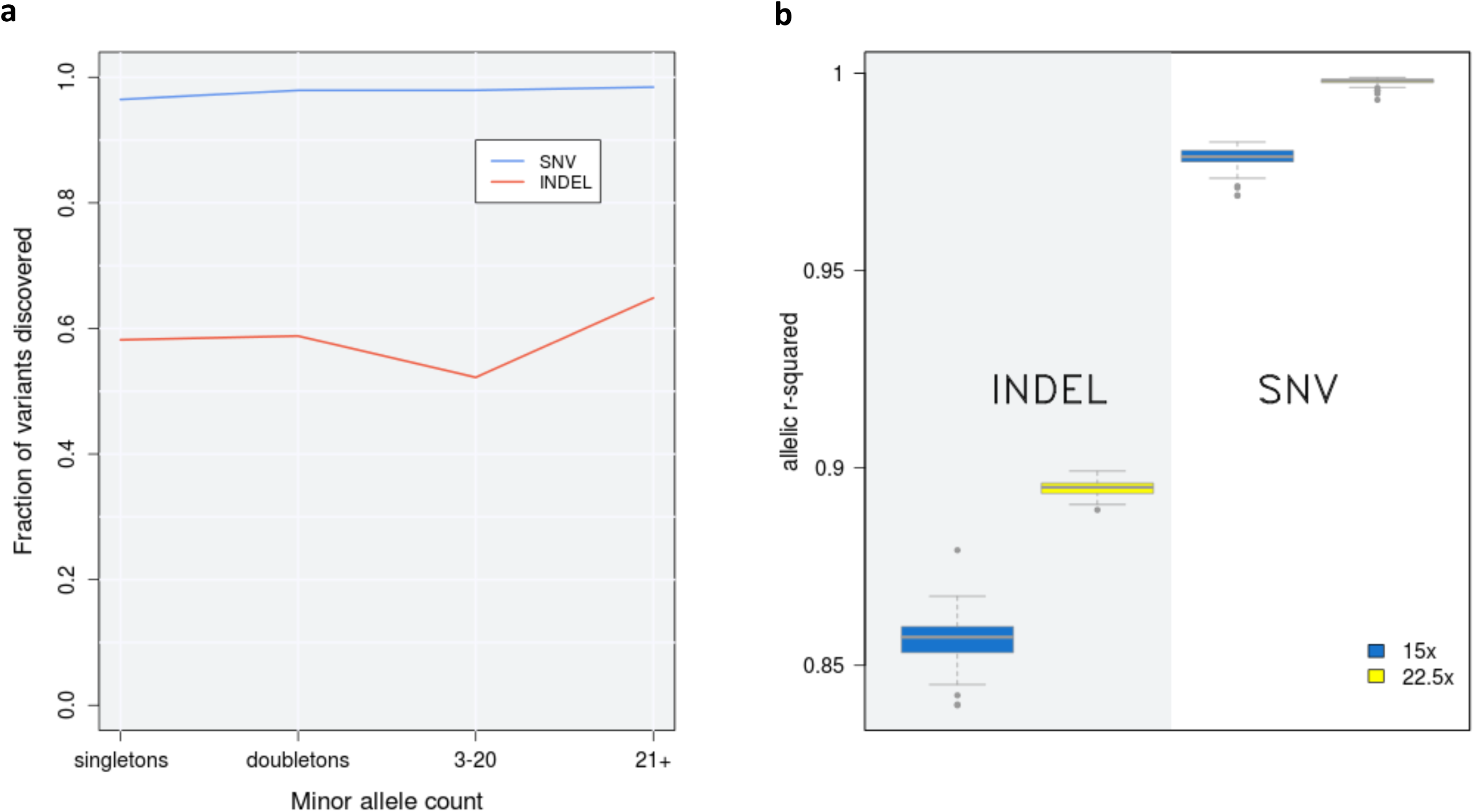
Variant discovery and quality at various sequencing depths in whole-genome sequencing data from 100 samples downsampled from 30x: (a) variant discovery rate in 22.5x; (b) allelic r^2^ for SNVs and INDELs in both 15x and 22.5x calls. INDEL: insertion/deletion. SNV: single nucleotide variant.

**Figure 2.**
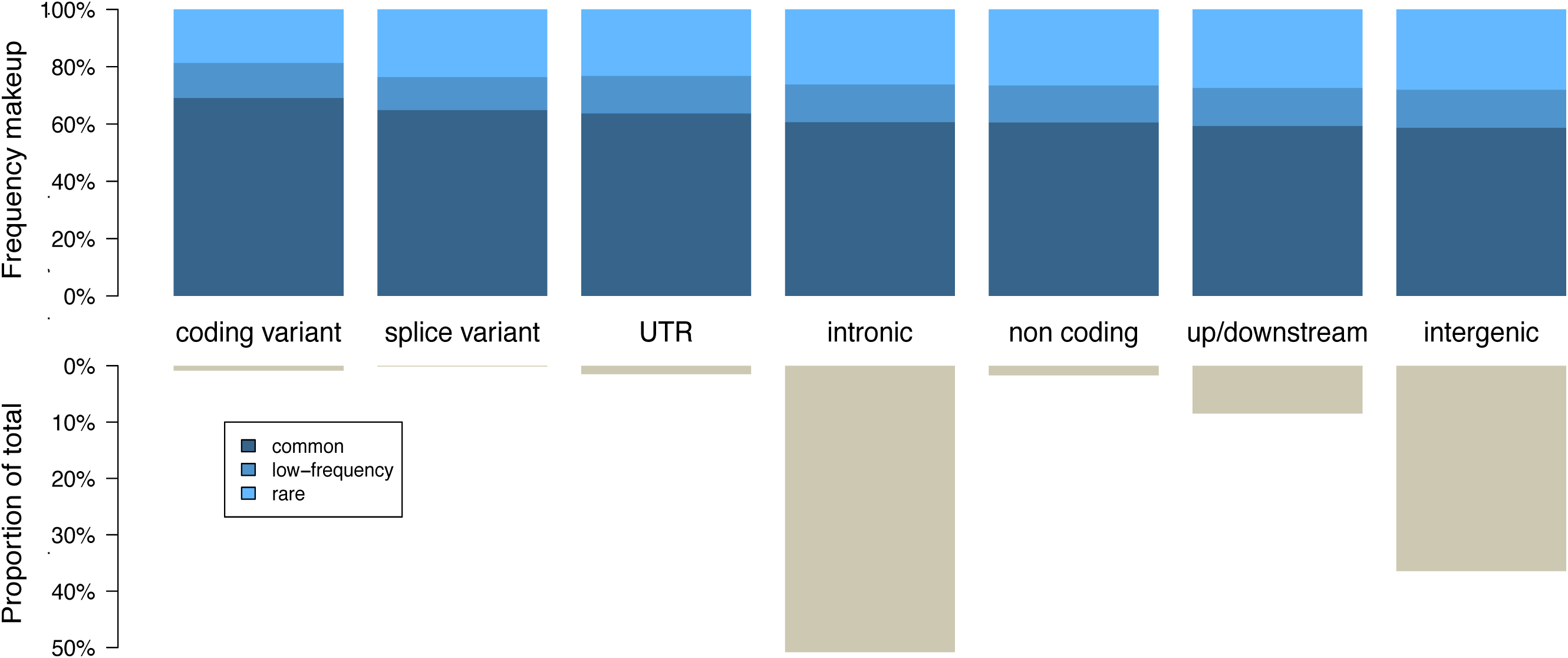
Variant count proportions and minor allele frequency bin by functional class. Functional classes are derived from the Ensembl VEP consequences as detailed in Supplementary Table 7, for 2 million randomly selected variants.

## RESULTS

### Effect of sequencing depth

Comparing whole genome sequencing at various depths ranging from 15x to 30x (Online Methods), we find that 96.4% of singletons, 97.9% of doubletons and 97.6% of variants called using 30x sequencing are recapitulated at 22.5x depth. Genotype accuracy (as measured by r^2^) is 99.7% for 22.5x depth and 98.5% for 15x depth, suggesting that increases between 15x and 30x translate into marginal improvements in both call rate and quality of very rare SNVs (Figure 1, Supplementary Figure 3 and Methods). We find that false discovery rates and genotype accuracy are substantially more dependent on sequencing depth for INDELs than for SNVs (Figure 1).

**Figure 3.**
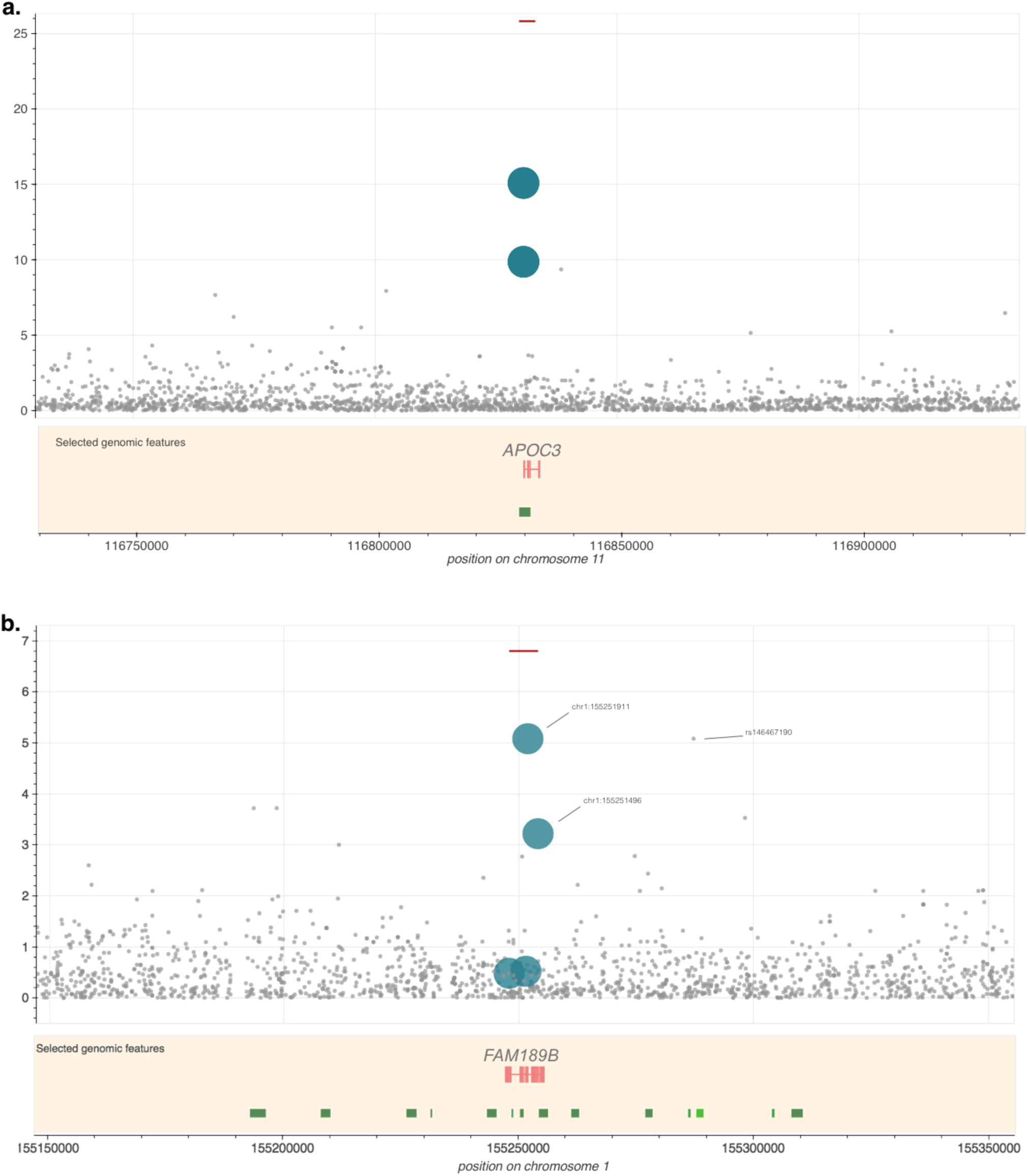
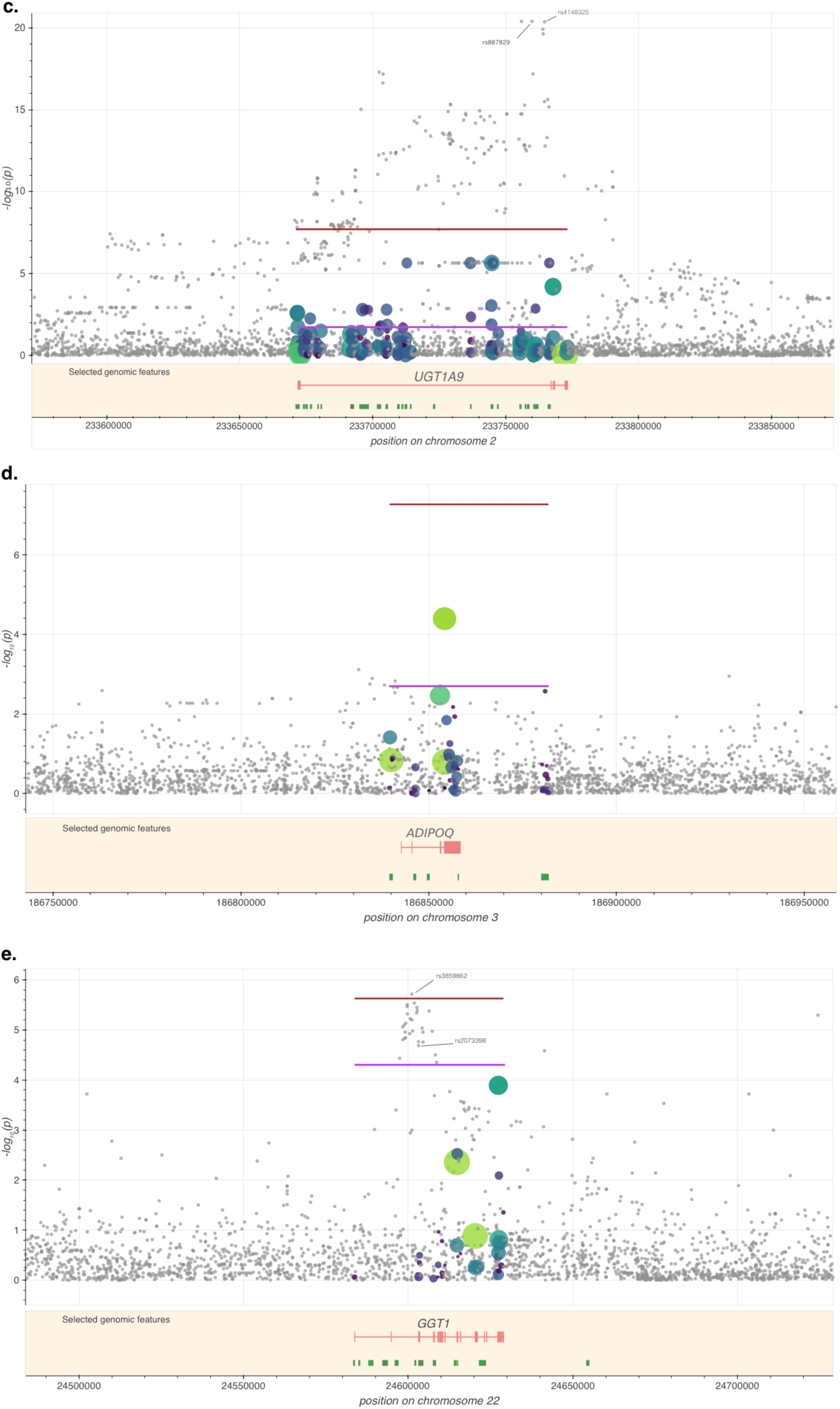
Regional association plots for burdens in *UGT1A9, ADIPOQ, FAM189B* and *GGT1*. Red lines denote the burden *P*-value and are drawn over the genomic region spanned by variants included in the test. Purple lines indicate the conditioned *P*-value for the variant described in the text, if applicable. Grey dots indicate single-point *P*-value for variants not included in the test. Coloured dots represent variants included in the test, with size and colour proportional to the score used in the most significant test, if applicable (Supplementary Tables 2 and 8). In the “Selected genomic features” panel, green bars below the gene denote regulatory regions associated with the gene.

### Landscape of sequence variation

Following quality control (QC), we call 24,163,896 non-monomorphic SNVs and INDELs, 97.9% of which are biallelic. 14,281,180 (60.31%) of the biallelic SNVs are rare (minor allele frequency [MAF]<0.01); 3,103,273 (13.1%) are low-frequency (MAF 0.01-0.05); and 6,292,726 (26.57%) are common (MAF>0.05). We call 8,294 non-monomorphic variants annotated as loss-of-function (LoF) with low-confidence (LC)^4^, and 438 variants annotated as LoF with high-confidence (HC) (Supplementary Figure 4). On average, each individual carries 405 (σ=19) LC LoF variants and 31 (σ=6) HC LoF variants, compared to 149 LoF variants per sample in a whole genome sequencing study of 2,636 Icelanders^5^. 0.6% and 1% of HC and LC LoF carrier genotypes are homozygous, respectively. INDELs are significantly more frequent among LoF variants, with 53.2% and 76% in the low- and high-confidence sets, respectively, compared to 13.5% genome-wide. We observe an enrichment of rare variants among the coding and splice variant categories (*P*=9.5×10^−16^) (Figure 2), and a lower rate of singletons compared to the general Greek population (*P*=10^−167^, one-sided empirical *P*-value) (Online Methods and Supplementary Figure 5), in keeping with the isolated nature of this Cretan population. Among the 5,102,175 novel biallelic variants (not present in gnomAD^6^ or Ensembl release 84^7^), 4,394,678 are SNVs, and the majority are rare (Supplementary Figure 6).

### Refinement of parameters for rare variant burden testing

We carried out genome-wide rare variant burden analyses for six medically-relevant traits: serum adiponectin, bilirubin, gamma-glutamyltransferase, low- and high-density lipoprotein, and triglyceride levels. As choice of genomic region, variant selection and weighting remain open questions for rare variant analysis, we benchmark 10 approaches using different regions of interest (exonic, exonic and regulatory, and regulatory only), variant inclusion and weighting methods (Online Methods; Supplementary Table 1). Overall, association statistics correlate highly within three distinct clusters (Supplementary Figure 7). Among exonic-only analyses, rare variant tests that only include unweighted high-consequence variants cluster separately from those in which variants are weighted according to their functionality scores. The third cluster encompasses all tests that include regulatory variants. Neither the variant weighting scheme nor the transformation used for adjusting the weights has a notable influence on the results.

### Rare variant burden discovery

In total, twenty burden signals exceed the study-wide significance threshold of 2.0×10^−7^ (Supplementary Figure 8), arising from four independent genes. Providing proof-of-principle, we identify association of a burden of loss-of-function variants with blood triglyceride and high-density lipoprotein levels in the *APOC3* gene (Figure 3.a, Supplementary Table 2)^2,8^. The strongest signal arises when the splice-donor variant rs138326449 (minor allele count (MAC)=38, minor allele frequency (MAF)=0.013) and the stop-gained variant rs76353203 (MAC=62, MAF=0.022) are included in the analysis (*P*=1.6×10^−26^). We replicate the association of a burden of rare coding *APOC3* variants with triglyceride levels in an independent dataset of 3,724 individuals with whole genome sequencing from the UK-based INTERVAL cohort^9^, in which we identify a burden of 25 exonic variants (*P* =3.1×10^−6^) (Supplementary Table 3). This is driven by rs138326449 and rs187628630, a rare 3’ UTR variant (MAF=0.008), with a two-variant burden *P*=9.0×10^−7^. rs138326449 is the only loss-of-function variant in *APOC3* present in this cohort, and is four times rarer than in MANOLIS (MAF_INTERVAL_=0.003 vs MAF_MANOLIS_=0.013).

We detect a new association of triglyceride levels with rare variants in the *FAM189B* gene (Figure 3.b, Supplementary Table 2). The burden association (*P*=1.56×10^−7^) is driven by two independent novel splice variants: chr1:155251911 G/A (human genome build 38, MAC=3, *P*=1.9×10^−6^) and chr1:155254079 C/G (MAC=2, *P*=1.9×10^−6^). In both cases, the minor allele is associated with increased triglyceride levels (β=2.59 σ=0.57 and β=2.40 σ=0.69, respectively). Both variants exhibit high quality scores (VQSLOD>19), high sequencing read depth (24x and 26.5x, respectively) and no missingness. A further novel splice region variant (chr1:155251496 T/C) and a stop gained variant (rs145265828), both singletons, were also included in the analysis; however their contribution to the burden is insignificant (burden *P*=2.26×10^−8^ when excluding them). We replicate evidence for a burden signal at *FAM189B* in the INTERVAL cohort (*P*=9.3×10^−3^) (Supplementary Table 3), which includes two stop gained variants with one driving the association: chr1:155250417 (rs749626426, MAC=2, β=1.96 σ=0.70, *P*=5.4×10^−3^). In keeping with the discovery dataset, the disruptive minor allele is associated with increased triglyceride levels. The two novel splice-region variants discovered in MANOLIS are not present in either the INTERVAL study or in a compendium of 123,136 exomes and 15,496 whole genomes assembled as part of the gnomAD project^6^. *FAM189B* has not been previously associated with blood lipid levels.

We find evidence of a low frequency and rare variant burden association with bilirubin levels in the *UGT1A9* gene (Figure 3.c, Supplementary Table 2). This association arises from the analyses including exonic and regulatory variants (*P*=1.9×10^−8^), and from the analyses including regulatory variants only (*P*=7.2×10^−8^). We find evidence for association in the exon plus regulatory region burden analysis in the INTERVAL replication cohort (*P*=1.7×10^−45^, Supplementary Table 3). A common variant in the first intron of *UGT1A9* (rs887829, MAF=0.28, β=0.426 σ=0.04, *P*=4.0×10^−21^ in the MANOLIS cohort) has previously been associated with bilirubin levels^10,11^. As expected, genotype correlation between rs887829 and each of the low-frequency and rare variants included in the burden is low (r_max_^2^=0.1). The rs887829 signal is not attenuated when conditioning on carrier status for the two main drivers of the burden (*P*_*conditional*_=4.5×10^−21^), or when conditioning on the number of rare alleles carried per individual (*P*_*conditional*_=4.0×10^−21^). The evidence for association with the rare variant burden in *UGT1A9* is substantially reduced when conditioned on rs887829 (*P*_*conditional*_=0.0146). Conversely, the two-variant signal for the two main burden drivers is attenuated from *P*=1.4×10^−7^ to *P*_*conditional*_=7.0×10^−3^ when conditioning on rs887829, indicating that it likely recapitulates part of a signal driven by a known common-variant association in the region.

We identify an association of adiponectin levels with low-frequency and rare variants in the *ADIPOQ* gene (Figure 3.d, Supplementary Table 2). The evidence for association is stronger for exonic and regulatory variants combined (*P*=4.2×10^−8^) than in either the regulatory-only (*P*=0.19) or exon-only (*P*=2.0×10^−6^) analyses, suggesting a genuine contribution of both classes of variants to the burden. Missense variant rs62625753 (MAF=0.031, *P*=4.0×10^−5^) contributes to the burden signal and is predicted to be damaging. The strength of association for the burden is reduced, but not entirely attenuated, when conditioned on rs62625753 (*P*_*conditional*_=8.9×10^−4^), indicating that it is not singly driven by this variant. rs35469083 (MAF=0.044) also contributes to the burden, and is an expression quantitative trait locus (eQTL) for *ADIPOQ* in visceral adipose tissue (minor allele associated with decreased gene expression). rs62625753 and rs35469083 have consistent directions of effect, with the minor alleles associated with reduced adiponectin levels, in keeping with their functional consequences on the gene (two-variant burden *P=*4.8×10^−7^). No common-variant signal for adiponectin levels is present in this region in our dataset. The burden signal remains significant upon conditioning on the genotypes of all variants with previous associations for adiponectin, type 2 diabetes or obesity that are polymorphic in MANOLIS (Supplementary Table 4).

In addition to the four genes that meet study-wide significance, we find gamma-glutamyltransferase levels to be suggestively associated with a burden of low frequency and rare exonic variants in the gamma-glutamyltransferase 1 (*GGT1*) gene (*P*=2.3×10^−6^) (Figure 3.e, Supplementary Table 2). A previously-reported, common-variant association is also present in an intron of this gene (rs3859862, MAF=0.46, *P*=1.9×10^−6^). The burden signal in *GGT1* is maintained when conditioning on rs3859862 (*P*_*conditional*_*=*5.1×10^−5^), suggesting that rare variants be independently contributing to this established association. Similarly, the single-point association at rs3859862 conditioned on carrier status for all rare variants included in the burden is not attenuated (*P*_*conditional*_=2.8×10^−5^), a result recapitulated by conditioning the same variant on the number of rare alleles carried per individual (*P*_*conditional*_=1.8×10^−5^), providing evidence for an independent rare variant signal at this locus.

### Signatures of selection

We surveyed the genomic loci with evidence of rare variant burden signals for signatures of recent positive selection in the MANOLIS cohort, using integrated haplotype scores (iHS)^12^. Notably, we find that 32% of the SNVs in *FAM189B* have an iHS score above 2, placing it in the top 5% of all genes analysed (96.7th percentile). This result is robust across several definitions of the genomic region representing the genes (95.6th-98.3th percentile) and to conditioning on gene length (94.6th percentile) (Supplementary Table 5).

## DISCUSSION

In this work, we have whole genome sequenced 1,457 individuals from the HELIC-MANOLIS cohort at an average depth of 22.5x. We describe the genomic variation landscape in this special population, discover 5.1 million novel variants, and perform rare variant burden testing across the entire genome for medically-relevant biochemical traits.

We empirically address several open whole genome sequencing study design and analysis questions. Through a downsampling approach, we demonstrate that it is possible to achieve near-perfect sensitivity and quality for rare variant calling and genotyping with half the depth, and at substantially lower cost, compared to 30x sequencing.

Defining the genomic regions in which to select variants, filtering strategies and variant weighting schemes constitute unresolved challenges in whole genome sequence-based studies. We find that association signal profiles of tests including regulatory region variants differ markedly from other scenarios, with some signals being driven by this variant class. Further, signal strength differs substantially between analyses that include high-severity consequence exonic variants only, and those in which all exonic variants are weighted according to their predicted consequence. We find that, as a rule, variant and functional unit selection, rather than weighting scheme, plays the largest role in association testing.

We identify a role for rare regulatory variants in the allelic architecture of complex traits. It is therefore important to leverage the whole genome sequence nature of the study data, and not to restrict analyses to coding variation only. We observe congruent directions of effect among regulatory and coding rare variants in burden signals that combine both classes of variation, for example across eQTL and damaging missense variants in the *ADIPOQ* gene that are together associated with adiponectin levels.

We discover replicating evidence for association of a rare variant burden with triglyceride levels at a locus not previously linked with the trait. *FAM189B* (Family With Sequence Similarity 189 Member B), also known as *COTE1* or *C1orf2*, codes for a membrane protein that is widely expressed, including in adult liver tissue^13^. Expression of *FAM189B* has been found to be correlated with endogenous SREBP-1 activation *in vitro*^14^. Sterol-regulatory element binding proteins (SREBPs) control the expression of genes involved in fatty acid and cholesterol biosynthesis, therefore indicating a mechanism by which *FAM189B* could be involved in lipid metabolism. We found *FAM189B* to display an elevated fraction of SNVs with |iHS|>2, a potential signature of recent positive selection at this genomic locus. This is particularly interesting in the context of this population, which has a high animal fat content diet^3^, and for which loss of function variants in *APOC3* have risen in frequency compared to the general population and confer a cardioprotective effect^15,16^.

We replicate the *FAM189B* association in an independent dataset with deep whole genome sequence data, in which the disruptive rare alleles are also associated with the same trait in the same direction. Across the board, we replicate all burden signals for which replication cohort trait measurements are available. We find that allelic heterogeneity is prevalent, partly due to the rare nature of the variants contributing to the burdens, and partly due to the distinct population genetics characteristics of the discovery and replication sets. These findings have important consequences for defining replication in sequence-based studies of rare variants, and highlight the importance of defining replication at the locus level rather than the variant level for burden signals.

We demonstrate pervasive allelic heterogeneity at complex trait loci, and identify exonic and regulatory rare variant associations at established signals. We find multiple instances of burden signals that remain independent of colocalising common variant signals, and one instance of burden signal attenuation when conditioning on the established common variant association. Within the power constraints of the study, we do not find evidence for synthetic association at established signals, i.e. there is no evidence for multiple rare variants at a locus accounting for a common variant association.

The discovery of rare variant burden associations with a modest sample size has been made possible due to the special population genetics characteristics of the isolated cohort under study. Rare variant signals, such as the ones discovered in *APOC3* and *FAM189B* in MANOLIS, are driven by variants with severe consequences that are rarer or absent in cosmopolitan populations. This demonstrates that the well-rehearsed power gains conferred by isolated cohorts in genome-wide association studies^17^ extend to whole genome sequence-based rare variant association designs.

Our findings indicate that deep whole genome sequencing at scale will be required to enable exhaustive description of the rare variant burden landscape in a population. For example, in the case of the *FAM189B* signal, low-depth sequencing (1x depth) of 1,239 MANOLIS samples^18^ misses one of the two burden-driving variants (chr1:155251911, MAC=3). Similarly, genome-wide genotyping coupled to dense imputation of the same samples does not capture the variants driving the burden signal identified here through deep whole genome sequencing^19^.

Our findings provide evidence for a role of low-frequency and rare, regulatory and coding variants in complex traits, and highlight the complex nature of locus-specific architecture at established and newly emerging signals. We anticipate that larger-scale, cohort-wide, deep whole genome sequencing initiatives will substantially further contribute to our understanding of the genetic underpinning of complex traits.

## Acknowledgements

HELIC-MANOLIS study: We thank the residents of the Mylopotamos villages for taking part. The MANOLIS study is dedicated to the memory of Manolis Giannakakis, 1978-2010. This work was funded by the Wellcome Trust [098051] and the European Research Council [ERC-2011-StG 280559-SEPI].

INTERVAL study: Participants in the INTERVAL randomised controlled trial were recruited with the active collaboration of NHS Blood and Transplant England (www.nhsbt.nhs.uk), which has supported field work and other elements of the trial. DNA extraction and genotyping was funded by the National Institute of Health Research (NIHR), the NIHR BioResource (http://bioresource.nihr.ac.uk/) and the NIHR Cambridge Biomedical Research Centre (www.cambridge-brc.org.uk). The academic coordinating centre for INTERVAL was supported by core funding from: NIHR Blood and Transplant Research Unit in Donor Health and Genomics, UK Medical Research Council (G0800270), British Heart Foundation (SP/09/002), and NIHR Research Cambridge Biomedical Research Centre. A complete list of the investigators and contributors to the INTERVAL trial is provided in reference^20^. This report is independent research by the National Institute for Health Research. The views expressed in this publication are those of the author(s) and not necessarily those of the NHS, the National Institute for Health Research or the Department of Health. This work was undertaken by Cambridge who received funding from the NHSBT; the views expressed in this publication are those of the authors and not necessarily those of the NHSBT.

TEENAGE study: The TEENAGE study has been supported by the Wellcome Trust (098051), European Union (European Social Fund—ESF) and Greek national funds through the Operational Program “Education and Lifelong Learning” of the National Strategic Reference Framework (NSRF)—Research Funding Program: Heracleitus II, Investing in knowledge society through the European Social Fund.

The GATK3 program was made available through the generosity of the Medical and Population Genetics program at the Broad Institute, Inc. We acknowledge Giuseppe Matullo’s contribution as EC’s PhD supervisor.

## Data access

Sequencing data are available at the European Genome-Phenome Archive under accession numbers EGAS00001001207 for MANOLIS, EGAS00001000988 for TEENAGE, and EGAS00001001355, EGAS00001002461 and EGAS00001002787 for INTERVAL. Code for the burden association pipeline and the burden visualisation tool is available at https://github.com/wtsi-team144/.

## Author Contributions

Sample collection and phenotyping: AEF, IN, ET, JD, GD, EZ

Sequencing Quality Control: AG, DS, KH, EC

Study design: AG, DS, KK, TB, EVRA, EZ

Association analyses: AG, DS, LS

Software development: AG, DS

Bioinformatics: AG, DS

Selection analysis: RW, IM

Phenotype data management: KK, GM, BK, NWR, AB

Downsampling/depth analysis: AG, MP

Replication cohort analyses: AG, KousikK, KW

Manuscript writing: AG, RW, IM, IB, EZ

Project supervision: EZ

## ONLINE METHODS

### Sequencing

For MANOLIS, genomic DNA (500 ng) from 1,482 samples was sheared to a median insert size of 500 bp and subjected to standard Illumina paired-end DNA library construction. Adapter-ligated libraries were amplified by 6 cycles of PCR and subjected to DNA sequencing using the HiSeqX platform (Illumina) according to manufacturer’s instructions. For TEENAGE, one hundred samples from the general Greek population were sequenced, as well as the Genome in a Bottle NA12878 sample. Sample identity checks were performed using Fluidigm and aliquots prepared. These aliquots underwent library preparation using the standard HiSeqX method. Size selection was performed to target 350 base pairs. Sequencing was performed on the Sanger Institute’s Illumina HiSeqX plat-form with a target depth of 30x and PhiX spike-in.

### Evaluation of sequencing accuracy at various depths

Reads from the NA12878 were downsampled to several read depths (from 5x to 30x) using the -s option of samtools view, aligned and processed through GATK Variant Quality Score Recalibrator. They were then compared to Genome in a Bottle (GIAB) 0.2 calls to extract the true positive rate (Supplementary Figure 2). At 22.5x, true positive rates are 98% for SNVs and 76% for INDELs.

### Comparison with the general Greek population

We compared singleton and doubleton counts in MANOLIS to 100 samples from the Greek general population (TEENAGE study), for which an identical sequencing protocol was used. The average depth in the TEENAGE study was 32.1x. We downsampled the individual BAMs to 22.5x and 15x using the -s option of samtools based on the average depth of the TEENAGE dataset, then performed variant calling using GATK HaplotypeCaller v3.3 (https://github.com/mp15/af_analysis) and filtering using GATK Variant Quality Score Recalibrator. The downsampled and original datasets were then compared using bcftools stats to extract allelic r-squared (Figure 1.b.). For the 22.5x dataset, we compared variant overlap with bcftools isec (Figure 1.a. and Supplementary Figure 3). We randomly drew 1,000 sets of 100 samples from the MANOLIS study and counted singletons and doubletons. We used these data to build a distribution and compute empirical quantiles for the singleton and doubleton counts calculated in TEENAGE. We counted 270,916 singletons and 61,690 doubletons in TEENAGE, compared with a median of 179,100 (p=1.4×10^−94^) and 75,280 (*P*=3.0×10^−19^), respectively, in MANOLIS (Supplementary Figure 5).

### Variant calling

Basecall files for each lane were transformed into unmapped BAMs using Illumina2BAM, marking adaptor contamination and decoding barcodes for removal into BAM tags. PhiX control reads were mapped using BWA Backtrack and were used to remove spatial artefacts. Reads were converted to FASTQ and aligned using BWA MEM 0.7.8 to the 1000 Genomes hs37d5 (for NA12878) and hg38 (GRCh38) with decoys (HS38DH) (for TEENAGE) references. The alignment was then merged into the master sample BAM file using Illumina2BAM MergeAlign. PCR and optical duplicates are marked using biobambam markduplicates and the files were archived in CRAM format.

Per-lane CRAMs were retrieved and reads pooled on a per-sample basis across all lanes to produce library CRAMs; these were each divided in 200 chunks for parallelism. GVCFs were generated using HaplotypeCaller v.3.5 from the Genome Analysis Toolkit (GATK) for each chunk. All chunks were then merged at sample level, samples were then further combined in batches of 150 samples using GATK CombineGVCFs v.3.5. Variant calling was then performed on each batch using GATK GenotypeGVCFs v.3.5. The resulting variant callsets were then merged across all batches into a cohort-wide VCF file using bcftools concat.

### Quality control

Variant-level QC was performed using the Variant Quality Score Recalibration tool (VQSR) from the Genome Analysis Toolkit (GATK) v. 3.5-0-g36282e4 ^21^, using a tranche threshold of 99.4% for SNPs, which provided an estimate false positive rate of 6%, and a true positive rate of 95%. For INDELs, we used the recommended threshold of 1%. For sample-level QC, we made extensive use of a previously described ^19^ GWAS dataset in 1175 overlapping samples. Four individuals failed sex checks, 8 samples had low concordance 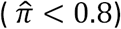 with chip data, 11 samples were duplicates, and 12 samples displayed traces of contamination (Freemix score from the verifyBamID suite ^22^ >5%). In case of sample duplicates, the sample with highest quality metrics (depth, freemix and chipmix score) was kept. As contamination and sex mismatches were correlated, a total of 25 individuals were excluded (N=1,457). No further samples were excluded based on depth, heterozygosity, transition/transversion (Ti/Tv) rate, missingness or ethnicity. No rare or low-frequency variant (MAF<5%) was excluded based on the Hardy-Weinberg equilibrium test at *P*=1.0×10^−5^. We filtered out 14% of variants with call rates <99%.

### Genetic relatedness matrix

Several methods are available to estimate the genetic relatedness present in isolated cohorts such as HELIC-MANOLIS^23^. We compared methods proposed in GEMMA^24^, EMMAX^25^, KING^26^ and PLINK.^27^, and found that the kinship coefficients reported by each method were highly correlated, but on a different scale from each other (Supplementary Figure 9). For consistency with previous studies performed on the same samples, we calculated a genetic relatedness matrix using GEMMA ^24^ after filtering for MAF<0.05, missingness <1% and LD-based pruning. In addition, MONSTER requires self-kinship coefficients on the diagonal of the relatedness matrix, which we calculated using the 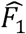 metric from PLINK 1.9. The matrix was then converted to the long format using the reshape2 R package.

### Association testing

Burden testing was performed using MONSTER^28^, a method that extends the SKAT-O^29^ model to account for relatedness and/or structure present in cohorts such as population isolates when testing for association. We ran burden testing across all genes defined in GENCODE v25 using 10 different conditions, i.e. combinations of regions of interest (coding regions only, coding and regulatory regions and regulatory regions only), variant filters (inclusion criteria based on severity of predicted consequence) and weighting schemes (Supplementary Table 1).

#### Examination of coding regions only

First, we extracted exonic coordinates for all protein-coding genes, which defines the region of interest for strictly exonic variants. These regions of interest were used in combination with 5 different variant filtering and weighting schemes. First, we included only variants predicted as high-confidence (HC) loss-of-function (LoF) by LOFTEE^4^ that reside in the exons of protein-coding genes (Supplementary Table 1: LOFTEE HC). As only 460 variants in 85 genes passed this inclusion criterion, we performed an additional analysis including 8,570 low-confidence (LC) loss-of-function variants spread across 1,727 genes (Supplementary Table 1: LOFTEE LC). Stop-gained and frameshift mutations were the largest contributors to both the LC and HC sets. However, the LC set also includes a large number of splice donor and splice acceptor variants (Supplementary Figure 4). We further performed an analysis with more relaxed inclusion criteria, including all exonic variants for which the Ensembl most severe consequence was more damaging than missense as predicted by the Variant Effect Predictor^30^ (Supplementary Table 1: Exon severe). We also employed Combined Annotation Dependent Depletion (CADD)^31^ scores, either to weigh all exonic variants (Supplementary Table 1: Exon CADD) or to filter out variants with CADD scores below the genome-wide median (Supplementary Table 1: Exon CADD median). Finally, we extended exon boundaries as defined above with 50 base pairs either side, to account for cases where potentially damaging variants occur on the edges of exons, as has been shown to happen for previously identified rare variant burdens^2^. These regions of interest were used in combination with one variant weighting scheme only (Exon+50 CADD).

#### Examination of regulatory and coding regions

We extracted regulatory regions (promoters, enhancers and transcription-factor binding sites) from Ensembl build 84^7^. We assigned regulatory regions to genes if they directly overlapped or if the regulatory region overlapped with an eQTL for the gene based on the GTEx database^32^. If an eQTL was reported for several genes, overlapping variants were assigned to all of them. We did not take tissue specificity into account. For selecting variants, we either used the coordinates of the regulatory features alone, or regulatory features plus the extended exons. We used Eigen, an aggregate score that combines information from multiple regulatory annotation tracks^33^, to weigh variants in all tests that include regulatory variants. In addition to raw Eigen scores, the authors also proposed EigenPC, a score derived from the first eigenvector of the correlation matrix of annotations. Both scores were available as is, or transformed using Phred-scaling, which maps a distribution’s support to [1, +∞], thereby guaranteeing inclusion and relative up-weighting of all variants. In the regulatory regions plus exon analyses we used both the raw Eigen scores, shifted by 1 unit to the right, with negative scores set to 0 +*ϵ* (Supplementary Table 1: Exon and regulatory Eigen), and the Phred-transformed Eigen and EigenPC scores (Supplementary Table 1: Exon and regulatory EigenPhred and EigenPCPhred). This transformation was a technical requirement as MONSTER could only read weights belonging to [0, +∞]. In the analyses containing the regulatory regions only, variants were weighted using the Phred-scaled Eigen scores (Supplementary Table 1: regulatory only EigenPhred) only.

Finally, we applied a MAF threshold of 0.05, a missingness threshold of 1% and a Hardy-Weinberg filter using a mid-p adjusted *P*-value^34^ threshold of 1.0×10^−5^ to all variants prior to testing. We only performed a test if at least two SNVs passed the inclusion criteria for a given condition.

### Establishing the significance threshold

We calculated 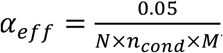, where > is the number of genes tested, *n*_*cond*_ is the effective number of inclusion and weighting criteria tested and *M* = 6 is the number of traits. For *n*_*comb*_, we plotted the correlation matrix of z-scores for all 10 analyses, and determined that the analyses using similar region definitions (exonic loss-of-function, exonic, exonic and regulatory variants) cluster together, reducing the effective number of analyses to 3 (Supplementary Figure 7). Although *N*=18,997 protein-coding genes are available in GENCODE V25, not all genes were tested in every condition. For example, for many genes only one variant might pass inclusion criteria in a high-confidence loss-of-function run, thereby excluding those genes from the analysis. A summary of the number of genes included in every analysis is presented in Supplementary Table 6. On average, *N*=13,854 genes are included, hence we define study-wide significance at *P*=2.0×10^−7^.

### Burden prioritisation and novelty

We applied stringent checks to test the validity of rare variant burden association signals. Every suggestively associated burden (arbitrarily defined as *P*≤5.0×10^−5^) was conditioned on the genotypes of the variant included in the burden set with the lowest single-point *P*-value. If the *P*-value dropped more than two orders of magnitude below the suggestive significance threshold (i.e. *P*≤5.0×10^−3^), the burden was excluded from downstream analyses. We examined burden signals using the plotburden software (https://github.com/wtsi-team144/plotburden) to assess variant functionality, single-point association *P*-values, LD structure, as well as prior associations in the region. When a prior association was found in the region, we considered a signal known when the *P*-value dropped below *P*=1.0×10^−4^ when conditioning on the genotypes of the existing signal.

### Replication

The INTERVAL randomized controlled trial is a large-scale study focusing on healthy blood donors^9^. Sequencing, variant calling and quality control was performed for 3,762 INTERVAL participants using the same protocol and pipeline as for the MANOLIS sequences. 38 samples were excluded on the basis of ethnicity, excessive relatedness (pi-hat>0.125), excess heterozygosity and contamination. VQSR thresholds of 99% and 90% for SNVs and INDELs, respectively, were applied to variant calls. Gamma-glutamyltransferase and adiponectin levels were not available in the INTERVAL replication cohort.

### Selection analysis

For the selection analysis we used the haplotype-based approach iHS ^12^. Previous studies have shown that an elevated fraction of SNVs with |iHS|>2 in a genomic region is a signature of selection^12,35^. Thus, to investigate if any of the lead genes identified in the burden tests have undergone positive selection, we focused on the fraction of SNVs with |iHS|>2. Specifically, we calculated this fraction for the four genes with study-wide significant association signals and assessed if they were elevated by comparing them to the empirical distribution for all genes.

#### Data sources

The primary data used for the selection analyses is the MANOLIS genotype data described above, which we phased using Beagle v.4.1 ^36^. We also used the ancestral allele annotations for each site in hg38 from ENSEMBL. Finally, we used the recombination map from UCSC, which was built on hg37 and lifted-over to hg38. From this map, we excluded 3140 sites to achieve a recombination map with monotonically increasing cM positions. Linear interpolation was subsequently used to produce cM positions for all sites not found on the map. For quality control, we combined the MANOLIS genotype data with genotype data from the 1000 Genomes phase 3^37^.

#### Filtering of individuals

Because close relatives can complicate and potentially bias analyses of signals of selection, we removed close relatives within the MANOLIS data. We used the same criteria as in ^1^, i.e. we used the --genome option in PLINK 1.9 to estimate PI-HAT and randomly excluded one individual from each pair of individuals with PI-HAT>0.2. These exclusions left 810 unrelated individuals from MANOLIS, on which we based the selection analysis.

#### Filtering of sites

We restricted our iHS analysis to known, common SNVs with ancestral allele annotations. Specifically, we excluded sites: not on the autosomes, with more than 2 alleles, with alleles that were not length 1 (INDEL-like), with MAF<0.05, without ancestral allele annotations, with HWE midp-value < 1e-30, not present in 1000 Genomes phase 3 vcf files, or that were outside a mappable region of the hg38 reference genome, defined as a GEM 100-mer score below 0.8 ^38^. These filtering steps resulted in 5,126,987 SNVs as input for iHS calculations.

#### iHS calculation

The iHS statistic was calculated with the hapbin program ^39^, using default parameters. The raw iHS statistic is sensitive to allele frequency, so SNVs were subsequently binned by derived allele count (82 equally-spaced bins) and the iHS statistic was normalized within each bin to have a mean of zero and a standard deviation of one, as suggested in ^12^. Finally, we examine the absolute value of the normalized iHS statistic to capture selection signals involving both derived and ancestral alleles. The iHS statistic for each site is affected by the extent of haplotypes its alleles exist on. Due to edge effects at chromosome ends and other gaps, we examine iHS values for 5,116,861 SNVs (99.8% of input sites).

#### Calculating fractions of SNVs with |iHS|>2 for a given gene

For each gene, we considered four distinct ways to define the genomic region representing the gene: 1) sites within exons, 2) sites within exons extended by 50bp or in regulatory elements, 3) sites within the region spanned by connecting all exons, and 4) sites within the region spanned by connecting all exons extended by 50bp and regulatory elements. For each gene and each of the four genic region definitions, we then extracted the SNVs with an iHS value and calculated the fraction of SNVs with normalized |iHS| above 2. When interpreting our results, we mainly focused on the results for the most inclusive definition, definition 4, as selection signatures tend to span fairly large genomic regions, but included the other definitions to be able to assess if this choice of definition markedly affected our results.

#### Comparison of the fraction of SNVs with |iHS|>2 between burden signal genes and all other genes

For each lead gene with a rare variant study-wide significant burden signal (*APOC3, UGT1A9, ADIPOQ*, and *FAM189B*), we compared its fraction of SNVs with |iHS|>2 to all other genes with at least 1 iHS value-bearing SNV, using each of the four different gene region definitions. Each comparison was quantified by the percentile of genes with a higher fraction of SNVs with |iHS|>2. *FAM189B* was the only of the four burden genes with a fraction |iHS|>2 above zero. For this gene, we also performed a comparison to the subset of genes with a similar number of SNVs with iHS values as *FAM189B* (defined as +/- 10% of the number of SNVs with iHS in *FAM189B*) to ensure the varying number of SNVs in the genes we compared *FAM189B* to did not drastically affect the percentiles. Note that with the gene definitions used some SNVs will be included in several genes and thus the data points in the empirical distribution used for comparison are not entirely independent.

