## Supplementary Materials for "Cohort-wide deep whole genome sequencing and the allelic architecture of complex traits"

**SUPPLEMENTARY MATERIAL**

**Supplementary Table 1**
Region definition, variant selection and weighting systems used to define testing conditions for burden analysis.

| Burden analysis condition | Weighting system | Criterion | Exons | Regulatory Regions |
|---|---|---|---|---|
| LOFTEE HC | none | Predicted LoF by LOFTEE with high confidence | yes | no |
| LOFTEE LC | none | Predicted LoF by LOFTEE, both high and low confidence | yes | no |
| Exon severe | none | Ensembl most severe consequence more severe than missense | yes | no |
| Exon CADD | CADD | none | yes | no |
| Exon CADD median | CADD | CADD>5.851 | yes | no |
| Exon+50 CADD | CADD | none | yes extended by 50bp | no |
| Exon+Regulatory Eigen | Eigen (raw score + 1) | Eigen>0 | yes extended by 50bp | yes |
| Exon+Regulatory EigenPhred | Phred-transformed Eigen | none | yes extended by 50bp | yes |
| Exon+Regulatory EigenPCPhred | Phred-transformed EigenPC | none | yes extended by 50bp | yes |
| Regulatory only EigenPhred | Phred-transformed Eigen | none | no | yes |

**Supplementary Table 2**

**Variants included in the burdens of the selected associations.** Alleles: reference/non-reference alleles; Consequence: the most severe consequence predicted by Ensembl VEP; MAF: minor allele frequency; AC: non-reference allele count; Weight: the weight assigned to the variant in the most significant test as per Supplementary Figure 8; NA indicates that no weight was applied.

| Trait | Gene | Analysis | Variant ID (GRCh38) | rsID | Alleles | Consequence | MAF | AC | Weight | single-point *P*-value |
|---|---|---|---|---|---|---|---|---|---|---|
| Triglycerides | *APOC3* | LOFTEE LC | chr11:116830637 | rs76353203 | C/T | Stop gained | 0.0213 | 62 | NA | 8.33E-16 |
| Triglycerides | *APOC3* | LOFTEE LC | chr11:116830638 | rs138326449 | G/A | Splice donor | 0.0130 | 38 | NA | 1.39E-10 |
| Triglycerides | *FAM189B* | Exon severe | chr1:155251911 | - | G/A | Splice region | 0.0010 | 3 | NA | 8.27E-06 |
| Triglycerides | *FAM189B* | Exon severe | chr1:155254079 | - | C/G | Splice region | 0.0007 | 2 | NA | 6.04E-04 |
| Triglycerides | *FAM189B* | Exon severe | chr1:155251496 | - | T/C | Splice region | 0.0003 | 1 | NA | 3.01E-01 |
| Triglycerides | *FAM189B* | Exon severe | chr1:155247925 | rs145265828 | G/A | Stop gained | 0.0003 | 1 | NA | 3.30E-01 |
| Bilirubin | *UGT1A9* | Exon+reg. Eigen | chr2:233671124 | - | G/T | Intron | 0.0003 | 1 | 0.92 | 4.66E-01 |
| Bilirubin | *UGT1A9* | Exon+reg. Eigen | chr2:233671195 | rs183298272 | G/A | Intron | 0.0010 | 3 | 0.46 | 2.65E-01 |
| Bilirubin | *UGT1A9* | Exon+reg. Eigen | chr2:233671246 | rs3806598 | A/C | Intron | 0.0357 | 104 | 0.70 | 1.23E-01 |
| Bilirubin | *UGT1A9* | Exon+reg. Eigen | chr2:233671257 | rs17868321 | C/T | Intron | 0.0079 | 23 | 0.76 | 5.47E-01 |
| Bilirubin | *UGT1A9* | Exon+reg. Eigen | chr2:233671290 | rs17863775 | A/G | Intron | 0.0010 | 3 | 0.88 | 2.39E-01 |
| Bilirubin | *UGT1A9* | Exon+reg. Eigen | chr2:233671505 | - | A/G | Intron | 0.0007 | 2 | 0.91 | 2.20E-01 |
| Bilirubin | *UGT1A9* | Exon+reg. Eigen | chr2:233671659 | rs6714486 | T/A | Intron | 0.0285 | 83 | 1.14 | 2.50E-03 |
| Bilirubin | *UGT1A9* | Exon+reg. Eigen | chr2:233671719 | - | C/T | Intron | 0.0027 | 8 | 0.64 | 2.82E-01 |
| Bilirubin | *UGT1A9* | Exon+reg. Eigen | chr2:233671848 | rs17868322 | G/A | Intron | 0.0402 | 117 | 1.08 | 1.77E-02 |
| Bilirubin | *UGT1A9* | Exon+reg. Eigen | chr2:233672032 | rs72551330 | T/C | Missense | 0.0086 | 25 | 0.64 | 4.62E-01 |
| Bilirubin | *UGT1A9* | Exon+reg. Eigen | chr2:233672159 | rs41264153 | G/A | Synonymous | 0.0034 | 10 | 1.68 | 3.85E-01 |
| Bilirubin | *UGT1A9* | Exon+reg. Eigen | chr2:233672162 | - | T/A | Synonymous | 0.0021 | 6 | 0.83 | 4.76E-01 |
| Bilirubin | *UGT1A9* | Exon+reg. Eigen | chr2:233672434 | rs151238339 | T/C | Missense | 0.0158 | 46 | 0.37 | 6.40E-01 |
| Bilirubin | *UGT1A9* | Exon+reg. Eigen | chr2:233672684 | - | T/G | Missense | 0.0003 | 1 | 0.56 | 1.96E-01 |
| Bilirubin | *UGT1A9* | Exon+reg. Eigen | chr2:233672736 | rs376424202 | A/G | Missense | 0.0010 | 3 | 1.79 | 7.90E-01 |
| Bilirubin | *UGT1A9* | Exon+reg. Eigen | chr2:233672763 | rs775497112 | T/G | Missense | 0.0003 | 1 | 1.65 | 6.67E-01 |
| Bilirubin | *UGT1A9* | Exon+reg. Eigen | chr2:233673890 | rs556955067 | C/T | Intron | 0.0003 | 1 | 0.75 | 3.53E-01 |
| Bilirubin | *UGT1A9* | Exon+reg. Eigen | chr2:233674004 | rs17862858 | A/G | Intron | 0.0165 | 48 | 0.78 | 4.62E-02 |
| Bilirubin | *UGT1A9* | Exon+reg. Eigen | chr2:233674100 | - | A/C | Intron | 0.0007 | 2 | 0.82 | 1.36E-01 |
| Bilirubin | *UGT1A9* | Exon+reg. Eigen | chr2:233674141 | rs28969703 | G/A | Intron | 0.0007 | 2 | 0.82 | 5.39E-01 |
| Bilirubin | *UGT1A9* | Exon+reg. Eigen | chr2:233674345 | rs45539636 | T/C | Intron | 0.0003 | 1 | 0.88 | 1.18E-01 |
| Bilirubin | *UGT1A9* | Exon+reg. Eigen | chr2:233675200 | rs12615708 | G/A | Intron | 0.0357 | 104 | 0.41 | 1.23E-01 |
| Bilirubin | *UGT1A9* | Exon+reg. Eigen | chr2:233675453 | - | T/C | Intron | 0.0017 | 5 | 0.48 | 8.43E-01 |
| Bilirubin | *UGT1A9* | Exon+reg. Eigen | chr2:233675454 | - | G/A | Intron | 0.0017 | 5 | 0.30 | 8.43E-01 |
| Bilirubin | *UGT1A9* | Exon+reg. Eigen | chr2:233675462 | rs79913681 | C/G | Intron | 0.0319 | 93 | 0.60 | 5.68E-02 |
| Bilirubin | *UGT1A9* | Exon+reg. Eigen | chr2:233676479 | rs34707303 | C/T | Intron | 0.0055 | 16 | 0.68 | 3.58E-01 |
| Bilirubin | *UGT1A9* | Exon+reg. Eigen | chr2:233676496 | - | A/G | Intron | 0.0017 | 5 | 0.79 | 5.44E-03 |
| Bilirubin | *UGT1A9* | Exon+reg. Eigen | chr2:233676786 | rs74435166 | C/G | Intron | 0.0165 | 48 | 0.95 | 4.62E-02 |
| Bilirubin | *UGT1A9* | Exon+reg. Eigen | chr2:233679279 | rs150747338 | C/T | Intron | 0.0007 | 2 | 0.35 | 1.31E-01 |
| Bilirubin | *UGT1A9* | Exon+reg. Eigen | chr2:233679337 | - | A/G | Intron | 0.0003 | 1 | 0.63 | 6.39E-01 |
| Bilirubin | *UGT1A9* | Exon+reg. Eigen | chr2:233679436 | rs752875090 | C/G | Intron | 0.0007 | 2 | 0.28 | 9.78E-01 |
| Bilirubin | *UGT1A9* | Exon+reg. Eigen | chr2:233679589 | rs113117053 | C/T | Intron | 0.0003 | 1 | 0.73 | 2.37E-01 |
| Bilirubin | *UGT1A9* | Exon+reg. Eigen | chr2:233680491 | rs142850766 | A/G | Intron | 0.0086 | 25 | 0.99 | 3.07E-02 |
| Bilirubin | *UGT1A9* | Exon+reg. Eigen | chr2:233680543 | rs184094797 | C/G | Intron | 0.0010 | 3 | 0.78 | 3.42E-01 |
| Bilirubin | *UGT1A9* | Exon+reg. Eigen | chr2:233691765 | - | G/A | 5-prime UTR | 0.0003 | 1 | 1.48 | 3.28E-01 |
| Bilirubin | *UGT1A9* | Exon+reg. Eigen | chr2:233691866 | rs45464992 | A/G | 3-prime UTR | 0.0045 | 13 | 1.40 | 5.01E-02 |
| Bilirubin | *UGT1A9* | Exon+reg. Eigen | chr2:233692049 | rs17868327 | C/A | 3-prime UTR | 0.0165 | 48 | 0.82 | 4.62E-02 |
| Bilirubin | *UGT1A9* | Exon+reg. Eigen | chr2:233692258 | rs754957875 | T/C | 3-prime UTR | 0.0003 | 1 | 0.88 | 1.08E-01 |
| Bilirubin | *UGT1A9* | Exon+reg. Eigen | chr2:233692449 | rs45568235 | C/T | Intron | 0.0007 | 2 | 0.94 | 5.39E-01 |
| Bilirubin | *UGT1A9* | Exon+reg. Eigen | chr2:233695282 | - | T/C | Intron | 0.0062 | 18 | 0.34 | 9.78E-01 |
| Bilirubin | *UGT1A9* | Exon+reg. Eigen | chr2:233695283 | rs546580338 | T/A | Intron | 0.0062 | 18 | 0.31 | 9.78E-01 |
| Bilirubin | *UGT1A9* | Exon+reg. Eigen | chr2:233695287 | rs775200577 | A/G | Intron | 0.0062 | 18 | 0.66 | 9.78E-01 |
| Bilirubin | *UGT1A9* | Exon+reg. Eigen | chr2:233695314 | - | C/T | Intron | 0.0010 | 3 | 0.29 | 7.91E-02 |
| Bilirubin | *UGT1A9* | Exon+reg. Eigen | chr2:233695871 | rs182203911 | C/T | Intron | 0.0021 | 6 | 0.71 | 8.61E-01 |
| Bilirubin | *UGT1A9* | Exon+reg. Eigen | chr2:233695897 | - | G/A | Intron | 0.0021 | 6 | 0.71 | 2.90E-02 |
| Bilirubin | *UGT1A9* | Exon+reg. Eigen | chr2:233696195 | rs17864691 | C/T | Intron | 0.0278 | 81 | 0.91 | 1.57E-03 |
| Bilirubin | *UGT1A9* | Exon+reg. Eigen | chr2:233696527 | - | A/C | Intron | 0.0003 | 1 | 1.01 | 4.15E-01 |
| Bilirubin | *UGT1A9* | Exon+reg. Eigen | chr2:233696840 | rs17864692 | G/A | Intron | 0.0278 | 81 | 0.57 | 1.57E-03 |
| Bilirubin | *UGT1A9* | Exon+reg. Eigen | chr2:233697582 | rs139693283 | T/C | Intron | 0.0003 | 1 | 0.59 | 1.96E-01 |
| Bilirubin | *UGT1A9* | Exon+reg. Eigen | chr2:233698070 | - | G/A | Intron | 0.0014 | 4 | 0.71 | 5.30E-01 |
| Bilirubin | *UGT1A9* | Exon+reg. Eigen | chr2:233698359 | rs17863784 | G/A | Intron | 0.0278 | 81 | 0.56 | 1.57E-03 |
| Bilirubin | *UGT1A9* | Exon+reg. Eigen | chr2:233701900 | rs182385495 | C/A | Intron | 0.0206 | 60 | 0.83 | 2.52E-01 |
| Bilirubin | *UGT1A9* | Exon+reg. Eigen | chr2:233702215 | - | T/C | Intron | 0.0003 | 1 | 0.70 | 3.28E-02 |
| Bilirubin | *UGT1A9* | Exon+reg. Eigen | chr2:233702369 | rs116955810 | G/A | Intron | 0.0093 | 27 | 0.34 | 1.40E-02 |
| Bilirubin | *UGT1A9* | Exon+reg. Eigen | chr2:233702821 | rs769310431 | G/T | Intron | 0.0003 | 1 | 1.04 | 3.14E-01 |
| Bilirubin | *UGT1A9* | Exon+reg. Eigen | chr2:233702910 | rs78155632 | T/A | Intron | 0.0093 | 27 | 0.58 | 1.40E-02 |
| Bilirubin | *UGT1A9* | Exon+reg. Eigen | chr2:233704961 | - | C/T | Intron | 0.0003 | 1 | 1.00 | 3.28E-01 |
| Bilirubin | *UGT1A9* | Exon+reg. Eigen | chr2:233704995 | - | G/T | Intron | 0.0003 | 1 | 0.75 | 2.02E-01 |
| Bilirubin | *UGT1A9* | Exon+reg. Eigen | chr2:233705012 | rs544165304 | T/A | Intron | 0.0007 | 2 | 0.69 | 6.57E-01 |
| Bilirubin | *UGT1A9* | Exon+reg. Eigen | chr2:233705067 | rs141940353 | C/T | Intron | 0.0003 | 1 | 0.33 | 9.76E-01 |
| Bilirubin | *UGT1A9* | Exon+reg. Eigen | chr2:233705231 | rs77358763 | T/C | Intron | 0.0278 | 81 | 0.78 | 1.57E-03 |

| | | | | | | | | | | |
|---|---|---|---|---|---|---|---|---|---|---|
| Bilirubin | *UGT1A9* | Exon+reg. Eigen | chr2:233705440 | rs72986482 | G/A | Intron | 0.0405 | 118 | 0.18 | 3.10E-02 |
| Bilirubin | *UGT1A9* | Exon+reg. Eigen | chr2:233705445 | rs79916929 | G/A | Intron | 0.0093 | 27 | 0.87 | 1.40E-02 |
| Bilirubin | *UGT1A9* | Exon+reg. Eigen | chr2:233705461 | rs142417974 | A/C | Intron | 0.0007 | 2 | 0.75 | 5.39E-01 |
| Bilirubin | *UGT1A9* | Exon+reg. Eigen | chr2:233705512 | - | G/C | Intron | 0.0069 | 20 | 0.56 | 5.62E-01 |
| Bilirubin | *UGT1A9* | Exon+reg. Eigen | chr2:233705564 | rs529349858 | C/A | Intron | 0.0007 | 2 | 0.20 | 7.86E-02 |
| Bilirubin | *UGT1A9* | Exon+reg. Eigen | chr2:233709411 | - | A/T | Intron | 0.0007 | 2 | 0.84 | 7.32E-02 |
| Bilirubin | *UGT1A9* | Exon+reg. Eigen | chr2:233709576 | rs180747204 | A/C | Intron | 0.0045 | 13 | 0.73 | 8.98E-01 |
| Bilirubin | *UGT1A9* | Exon+reg. Eigen | chr2:233709577 | rs186174292 | A/T | Intron | 0.0045 | 13 | 0.78 | 8.98E-01 |
| Bilirubin | *UGT1A9* | Exon+reg. Eigen | chr2:233709733 | rs190992275 | T/C | Intron | 0.0117 | 34 | 0.78 | 5.23E-01 |
| Bilirubin | *UGT1A9* | Exon+reg. Eigen | chr2:233710022 | rs190818817 | G/A | Intron | 0.0007 | 2 | 0.71 | 1.31E-01 |
| Bilirubin | *UGT1A9* | Exon+reg. Eigen | chr2:233711088 | rs145698870 | A/G | Intron | 0.0003 | 1 | 0.52 | 6.67E-01 |
| Bilirubin | *UGT1A9* | Exon+reg. Eigen | chr2:233711092 | rs138181984 | G/T | Intron | 0.0003 | 1 | 0.79 | 6.67E-01 |
| Bilirubin | *UGT1A9* | Exon+reg. Eigen | chr2:233711384 | rs547291485 | G/C | Intron | 0.0069 | 20 | 0.87 | 5.21E-01 |
| Bilirubin | *UGT1A9* | Exon+reg. Eigen | chr2:233711404 | - | C/T | Intron | 0.0096 | 28 | 0.66 | 2.02E-02 |
| Bilirubin | *UGT1A9* | Exon+reg. Eigen | chr2:233712151 | - | A/G | Intron | 0.0017 | 5 | 0.84 | 1.60E-01 |
| Bilirubin | *UGT1A9* | Exon+reg. Eigen | chr2:233712177 | rs28898598 | C/T | Intron | 0.0007 | 2 | 0.93 | 5.39E-01 |
| Bilirubin | *UGT1A9* | Exon+reg. Eigen | chr2:233712193 | rs144263331 | C/A | Intron | 0.0003 | 1 | 0.20 | 2.34E-01 |
| Bilirubin | *UGT1A9* | Exon+reg. Eigen | chr2:233712285 | - | C/A | Intron | 0.0003 | 1 | 0.29 | 1.91E-01 |
| Bilirubin | *UGT1A9* | Exon+reg. Eigen | chr2:233712384 | rs45489204 | G/A | Intron | 0.0003 | 1 | 0.96 | 9.59E-01 |
| Bilirubin | *UGT1A9* | Exon+reg. Eigen | chr2:233712795 | rs551571775 | G/A | Intron | 0.0003 | 1 | 0.78 | 8.15E-02 |
| Bilirubin | *UGT1A9* | Exon+reg. Eigen | chr2:233712842 | - | C/T | Intron | 0.0254 | 74 | 0.68 | 2.23E-06 |
| Bilirubin | *UGT1A9* | Exon+reg. Eigen | chr2:233714147 | rs548246419 | C/T | Intron | 0.0007 | 2 | 1.01 | 5.93E-01 |
| Bilirubin | *UGT1A9* | Exon+reg. Eigen | chr2:233714200 | - | C/G | Intron | 0.0017 | 5 | 0.91 | 6.81E-01 |
| Bilirubin | *UGT1A9* | Exon+reg. Eigen | chr2:233736722 | rs149071654 | G/A | Intron | 0.0254 | 74 | 0.80 | 2.23E-06 |
| Bilirubin | *UGT1A9* | Exon+reg. Eigen | chr2:233736925 | rs188632907 | G/A | Intron | 0.0034 | 10 | 0.66 | 4.29E-03 |
| Bilirubin | *UGT1A9* | Exon+reg. Eigen | chr2:233736990 | - | A/C | Intron | 0.0003 | 1 | 0.37 | 1.24E-01 |
| Bilirubin | *UGT1A9* | Exon+reg. Eigen | chr2:233736992 | rs193257830 | G/T | Intron | 0.0010 | 3 | 0.66 | 6.40E-01 |
| Bilirubin | *UGT1A9* | Exon+reg. Eigen | chr2:233744624 | - | T/A | Intron | 0.0003 | 1 | 0.88 | 3.88E-01 |
| Bilirubin | *UGT1A9* | Exon+reg. Eigen | chr2:233744646 | rs28899468 | T/C | Intron | 0.0360 | 105 | 0.84 | 8.80E-04 |
| Bilirubin | *UGT1A9* | Exon+reg. Eigen | chr2:233744651 | - | A/T | Intron | 0.0106 | 31 | 0.41 | 2.18E-01 |
| Bilirubin | *UGT1A9* | Exon+reg. Eigen | chr2:233744657 | - | C/T | Intron | 0.0106 | 31 | 0.60 | 2.18E-01 |
| Bilirubin | *UGT1A9* | Exon+reg. Eigen | chr2:233744674 | - | A/C | Intron | 0.0031 | 9 | 0.10 | 3.56E-01 |
| Bilirubin | *UGT1A9* | Exon+reg. Eigen | chr2:233744700 | - | C/T | Intron | 0.0086 | 25 | 0.81 | 1.29E-02 |
| Bilirubin | *UGT1A9* | Exon+reg. Eigen | chr2:233744768 | rs116261344 | G/A | Intron | 0.0254 | 74 | 1.18 | 2.23E-06 |
| Bilirubin | *UGT1A9* | Exon+reg. Eigen | chr2:233744794 | rs145652010 | G/C | Intron | 0.0003 | 1 | 0.93 | 6.67E-01 |
| Bilirubin | *UGT1A9* | Exon+reg. Eigen | chr2:233744892 | rs28900378 | C/T | Intron | 0.0007 | 2 | 0.24 | 5.39E-01 |
| Bilirubin | *UGT1A9* | Exon+reg. Eigen | chr2:233745061 | rs115414076 | T/C | Intron | 0.0257 | 75 | 0.72 | 2.23E-06 |
| Bilirubin | *UGT1A9* | Exon+reg. Eigen | chr2:233745246 | rs148872010 | G/A | Intron | 0.0017 | 5 | 1.00 | 7.01E-01 |
| Bilirubin | *UGT1A9* | Exon+reg. Eigen | chr2:233747111 | - | T/C | Intron | 0.0003 | 1 | 0.89 | 2.55E-01 |
| Bilirubin | *UGT1A9* | Exon+reg. Eigen | chr2:233747199 | - | G/A | Intron | 0.0003 | 1 | 0.58 | 5.36E-01 |
| Bilirubin | *UGT1A9* | Exon+reg. Eigen | chr2:233747227 | rs138794928 | C/T | Non-coding transcript exon | 0.0065 | 19 | 0.76 | 4.43E-02 |
| Bilirubin | *UGT1A9* | Exon+reg. Eigen | chr2:233755356 | - | C/A | Non-coding transcript exon | 0.0027 | 8 | 0.75 | 2.82E-01 |
| Bilirubin | *UGT1A9* | Exon+reg. Eigen | chr2:233755391 | rs115647781 | C/G | Non-coding transcript exon | 0.0034 | 10 | 1.02 | 3.94E-01 |
| Bilirubin | *UGT1A9* | Exon+reg. Eigen | chr2:233755438 | rs192713031 | C/G | Non-coding transcript exon | 0.0003 | 1 | 1.08 | 6.67E-01 |
| Bilirubin | *UGT1A9* | Exon+reg. Eigen | chr2:233755508 | - | C/T | Non-coding transcript exon | 0.0072 | 21 | 1.15 | 3.47E-01 |
| Bilirubin | *UGT1A9* | Exon+reg. Eigen | chr2:233755509 | rs143373661 | C/G | Non-coding transcript exon | 0.0295 | 86 | 1.08 | 1.45E-01 |
| Bilirubin | *UGT1A9* | Exon+reg. Eigen | chr2:233755562 | rs185113714 | G/A | Non-coding transcript exon | 0.0010 | 3 | 0.56 | 3.19E-01 |
| Bilirubin | *UGT1A9* | Exon+reg. Eigen | chr2:233755690 | - | C/T | Non-coding transcript exon | 0.0007 | 2 | 1.29 | 1.56E-01 |
| Bilirubin | *UGT1A9* | Exon+reg. Eigen | chr2:233755723 | rs376520446 | G/A | Non-coding transcript exon | 0.0007 | 2 | 0.82 | 1.56E-01 |
| Bilirubin | *UGT1A9* | Exon+reg. Eigen | chr2:233755734 | - | G/C | Non-coding transcript exon | 0.0007 | 2 | 0.47 | 3.29E-02 |
| Bilirubin | *UGT1A9* | Exon+reg. Eigen | chr2:233755783 | rs142854795 | C/G | Non-coding transcript exon | 0.0093 | 27 | 0.81 | 1.29E-01 |
| Bilirubin | *UGT1A9* | Exon+reg. Eigen | chr2:233757439 | - | A/G | Intron | 0.0010 | 3 | 0.92 | 3.42E-01 |
| Bilirubin | *UGT1A9* | Exon+reg. Eigen | chr2:233757608 | - | A/G | Intron | 0.0027 | 8 | 1.17 | 1.02E-01 |
| Bilirubin | *UGT1A9* | Exon+reg. Eigen | chr2:233760498 | rs4148323 | G/A | Missense | 0.0003 | 1 | 0.92 | 9.76E-01 |
| Bilirubin | *UGT1A9* | Exon+reg. Eigen | chr2:233760827 | rs148755655 | A/G | Synonymous | 0.0034 | 10 | 1.44 | 6.24E-01 |
| Bilirubin | *UGT1A9* | Exon+reg. Eigen | chr2:233760961 | rs35003977 | T/G | Missense | 0.0031 | 9 | 0.10 | 3.56E-01 |
| Bilirubin | *UGT1A9* | Exon+reg. Eigen | chr2:233761103 | rs371726341 | G/A | Missense | 0.0089 | 26 | 0.35 | 3.47E-01 |
| Bilirubin | *UGT1A9* | Exon+reg. Eigen | chr2:233761173 | rs771182197 | T/G | Intron | 0.0010 | 3 | 1.24 | 2.25E-01 |
| Bilirubin | *UGT1A9* | Exon+reg. Eigen | chr2:233761235 | rs35041092 | C/T | Intron | 0.0021 | 6 | 1.38 | 3.31E-01 |
| Bilirubin | *UGT1A9* | Exon+reg. Eigen | chr2:233761240 | rs17868341 | C/T | Intron | 0.0275 | 80 | 0.60 | 1.34E-03 |
| Bilirubin | *UGT1A9* | Exon+reg. Eigen | chr2:233761493 | rs34652311 | C/T | Intron | 0.0007 | 2 | 1.25 | 5.39E-01 |
| Bilirubin | *UGT1A9* | Exon+reg. Eigen | chr2:233761914 | rs28900396 | T/C | Intron | 0.0007 | 2 | 0.98 | 5.39E-01 |
| Bilirubin | *UGT1A9* | Exon+reg. Eigen | chr2:233761943 | rs542276464 | G/A | Intron | 0.0003 | 1 | 1.18 | 4.79E-01 |
| Bilirubin | *UGT1A9* | Exon+reg. Eigen | chr2:233761967 | - | A/T | Intron | 0.0003 | 1 | 0.71 | 4.00E-01 |
| Bilirubin | *UGT1A9* | Exon+reg. Eigen | chr2:233762010 | - | G/A | Intron | 0.0007 | 2 | 0.68 | 6.14E-01 |
| Bilirubin | *UGT1A9* | Exon+reg. Eigen | chr2:233766235 | - | T/C | Intron | 0.0031 | 9 | 0.70 | 7.51E-01 |
| Bilirubin | *UGT1A9* | Exon+reg. Eigen | chr2:233766327 | rs373926451 | G/A | Intron | 0.0017 | 5 | 0.41 | 1.60E-01 |
| Bilirubin | *UGT1A9* | Exon+reg. Eigen | chr2:233766390 | rs11679312 | C/T | Intron | 0.0254 | 74 | 0.66 | 2.23E-06 |
| Bilirubin | *UGT1A9* | Exon+reg. Eigen | chr2:233766403 | - | C/T | Intron | 0.0010 | 3 | 0.70 | 9.05E-01 |
| Bilirubin | *UGT1A9* | Exon+reg. Eigen | chr2:233766495 | - | C/T | Intron | 0.0010 | 3 | 0.47 | 5.12E-01 |
| Bilirubin | *UGT1A9* | Exon+reg. Eigen | chr2:233766771 | rs71528513 | C/T | Intron | 0.0003 | 1 | 0.99 | 4.04E-01 |
| Bilirubin | *UGT1A9* | Exon+reg. Eigen | chr2:233767183 | rs34650714 | C/T | Intron | 0.0003 | 1 | 0.77 | 6.67E-01 |
| Bilirubin | *UGT1A9* | Exon+reg. Eigen | chr2:233767812 | rs12471326 | T/C | Intron | 0.0226 | 66 | 1.28 | 6.26E-05 |
| Bilirubin | *UGT1A9* | Exon+reg. Eigen | chr2:233767947 | rs61764033 | C/T | Intron | 0.0010 | 3 | 1.02 | 7.93E-02 |
| Bilirubin | *UGT1A9* | Exon+reg. Eigen | chr2:233772367 | rs769720139 | C/T | Synonymous | 0.0003 | 1 | 2.14 | 9.09E-01 |
| Bilirubin | *UGT1A9* | Exon+reg. Eigen | chr2:233773050 | - | T/G | 3-prime UTR | 0.0003 | 1 | 1.14 | 6.41E-01 |
| Bilirubin | *UGT1A9* | Exon+reg. Eigen | chr2:233773233 | - | G/A | 3-prime UTR | 0.0038 | 11 | 1.08 | 3.16E-01 |
| Adiponectin | *ADIPOQ* | Exon+reg. Eigen | chr3:186839605 | - | G/T | Upstream gene | 0.0003 | 1 | 0.69 | 7.23E-01 |
| Adiponectin | *ADIPOQ* | Exon+reg. Eigen | chr3:186839708 | - | T/C | Upstream gene | 0.0014 | 4 | 1.24 | 3.87E-02 |
| Adiponectin | *ADIPOQ* | Exon+reg. Eigen | chr3:186840083 | rs76071583 | A/G | Upstream gene | 0.0065 | 19 | 2.17 | 1.46E-01 |
| Adiponectin | *ADIPOQ* | Exon+reg. Eigen | chr3:186840236 | - | G/T | Upstream gene | 0.0075 | 22 | 0.44 | 1.45E-01 |

| | | | | | | | | | | |
|---|---|---|---|---|---|---|---|---|---|---|
| Adiponectin | *ADIPOQ* | Exon+reg. Eigen | chr3:186840399 | - | G/A | Upstream gene | 0.0003 | 1 | 0.74 | 1.30E-01 |
| Adiponectin | *ADIPOQ* | Exon+reg. Eigen | chr3:186845559 | - | A/G | Splice region | 0.0003 | 1 | 0.73 | 9.81E-01 |
| Adiponectin | *ADIPOQ* | Exon+reg. Eigen | chr3:186846476 | - | C/T | Intron | 0.0027 | 8 | 0.71 | 7.56E-01 |
| Adiponectin | *ADIPOQ* | Exon+reg. Eigen | chr3:186846494 | rs146842550 | A/C | Intron | 0.0024 | 7 | 0.94 | 7.56E-01 |
| Adiponectin | *ADIPOQ* | Exon+reg. Eigen | chr3:186846500 | - | C/A | Intron | 0.0003 | 1 | 0.93 | 2.19E-01 |
| Adiponectin | *ADIPOQ* | Exon+reg. Eigen | chr3:186846505 | rs140645649 | A/G | Intron | 0.0007 | 2 | 0.88 | 9.98E-01 |
| Adiponectin | *ADIPOQ* | Exon+reg. Eigen | chr3:186850096 | - | C/T | Intron | 0.0010 | 3 | 0.35 | 8.46E-01 |
| Adiponectin | *ADIPOQ* | Exon+reg. Eigen | chr3:186850184 | rs148752829 | C/T | Intron | 0.0003 | 1 | 0.22 | 8.46E-01 |
| Adiponectin | *ADIPOQ* | Exon+reg. Eigen | chr3:186853027 | rs17366653 | T/C | Non-coding transcript exon | 0.0175 | 51 | 1.69 | 3.44E-03 |
| Adiponectin | *ADIPOQ* | Exon+reg. Eigen | chr3:186854237 | rs62625753 | G/A | Missense | 0.0302 | 88 | 2.01 | 4.02E-05 |
| Adiponectin | *ADIPOQ* | Exon+reg. Eigen | chr3:186854300 | rs17366743 | T/C | Missense | 0.0223 | 65 | 0.49 | 7.22E-01 |
| Adiponectin | *ADIPOQ* | Exon+reg. Eigen | chr3:186854365 | rs139742770 | T/C | Synonymous | 0.0010 | 3 | 2.31 | 1.61E-01 |
| Adiponectin | *ADIPOQ* | Exon+reg. Eigen | chr3:186854769 | rs4068 | C/T | 3-prime UTR | 0.0003 | 1 | 1.05 | 1.43E-02 |
| Adiponectin | *ADIPOQ* | Exon+reg. Eigen | chr3:186855179 | rs201246459 | G/T | 3-prime UTR | 0.0038 | 11 | 0.95 | 1.21E-01 |
| Adiponectin | *ADIPOQ* | Exon+reg. Eigen | chr3:186855400 | rs6444174 | C/T | 3-prime UTR | 0.0106 | 2883 | 1.11 | 1.02E-01 |
| Adiponectin | *ADIPOQ* | Exon+reg. Eigen | chr3:186855646 | rs6796106 | G/A | 3-prime UTR | 0.0017 | 5 | 0.84 | 5.55E-02 |
| Adiponectin | *ADIPOQ* | Exon+reg. Eigen | chr3:186855700 | rs201119230 | C/T | 3-prime UTR | 0.0007 | 2 | 0.67 | 4.67E-01 |
| Adiponectin | *ADIPOQ* | Exon+reg. Eigen | chr3:186856163 | rs111856964 | G/A | 3-prime UTR | 0.0003 | 1 | 1.16 | 2.16E-01 |
| Adiponectin | *ADIPOQ* | Exon+reg. Eigen | chr3:186856493 | rs35469083 | C/T | 3-prime UTR | 0.0439 | 128 | 0.60 | 6.68E-03 |
| Adiponectin | *ADIPOQ* | Exon+reg. Eigen | chr3:186856749 | - | G/A | 3-prime UTR | 0.0017 | 5 | 1.07 | 8.00E-01 |
| Adiponectin | *ADIPOQ* | Exon+reg. Eigen | chr3:186856847 | - | A/G | 3-prime UTR | 0.0010 | 3 | 0.63 | 7.05E-01 |
| Adiponectin | *ADIPOQ* | Exon+reg. Eigen | chr3:186856936 | rs200028669 | C/T | 3-prime UTR | 0.0017 | 5 | 0.67 | 1.17E-02 |
| Adiponectin | *ADIPOQ* | Exon+reg. Eigen | chr3:186856958 | rs149814403 | C/T | 3-prime UTR | 0.0021 | 6 | 0.92 | 2.08E-01 |
| Adiponectin | *ADIPOQ* | Exon+reg. Eigen | chr3:186857194 | rs200077203 | C/T | 3-prime UTR | 0.0199 | 58 | 1.10 | 8.63E-01 |
| Adiponectin | *ADIPOQ* | Exon+reg. Eigen | chr3:186857286 | rs200360323 | G/C | 3-prime UTR | 0.0021 | 6 | 0.87 | 5.08E-01 |
| Adiponectin | *ADIPOQ* | Exon+reg. Eigen | chr3:186857470 | - | C/G | 3-prime UTR | 0.0007 | 2 | 1.07 | 3.79E-01 |
| Adiponectin | *ADIPOQ* | Exon+reg. Eigen | chr3:186857641 | rs73187705 | G/A | 3-prime UTR | 0.0024 | 7 | 1.11 | 1.50E-01 |
| Adiponectin | *ADIPOQ* | Exon+reg. Eigen | chr3:186857952 | rs188684283 | T/C | 3-prime UTR | 0.0106 | 31 | 0.34 | 2.45E-01 |
| Adiponectin | *ADIPOQ* | Exon+reg. Eigen | chr3:186880230 | rs80037191 | G/T | Intergenic | 0.0443 | 129 | 0.62 | 1.86E-01 |
| Adiponectin | *ADIPOQ* | Exon+reg. Eigen | chr3:186880272 | - | T/C | Intergenic | 0.0003 | 1 | 0.70 | 8.47E-01 |
| Adiponectin | *ADIPOQ* | Exon+reg. Eigen | chr3:186880565 | rs79680763 | C/T | Intergenic | 0.0048 | 14 | 0.72 | 7.98E-01 |
| Adiponectin | *ADIPOQ* | Exon+reg. Eigen | chr3:186880738 | rs182158541 | C/T | Intergenic | 0.0072 | 21 | 0.90 | 7.87E-01 |
| Adiponectin | *ADIPOQ* | Exon+reg. Eigen | chr3:186881003 | rs74398340 | C/A | Intergenic | 0.0213 | 62 | 0.66 | 2.68E-03 |
| Adiponectin | *ADIPOQ* | Exon+reg. Eigen | chr3:186881182 | rs116337242 | T/A | Intergenic | 0.0024 | 7 | 0.81 | 3.38E-01 |
| Adiponectin | *ADIPOQ* | Exon+reg. Eigen | chr3:186881423 | rs140486912 | T/C | Intergenic | 0.0460 | 134 | 0.48 | 2.00E-01 |
| Adiponectin | *ADIPOQ* | Exon+reg. Eigen | chr3:186881714 | rs535166771 | C/T | Intergenic | 0.0281 | 82 | 0.71 | 4.26E-01 |
| Adiponectin | *ADIPOQ* | Exon+reg. Eigen | chr3:186881715 | - | G/A | Intergenic | 0.0058 | 17 | 0.67 | 8.94E-01 |
| Adiponectin | *ADIPOQ* | Exon+reg. Eigen | chr3:186881882 | - | G/C | Intergenic | 0.0014 | 4 | 0.97 | 8.55E-01 |
| γ-glutamyltransferase | *GGT1* | Exon CADD | chr22:24583771 | rs774266253 | G/A | 5-prime UTR | 0.0014 | 4 | 2.09 | 8.68E-01 |
| γ-glutamyltransferase | *GGT1* | Exon CADD | chr22:24603184 | rs554233625 | C/T | 5-prime UTR | 0.0124 | 36 | 6.00 | 8.51E-01 |
| γ-glutamyltransferase | *GGT1* | Exon CADD | chr22:24603412 | - | G/C | 5-prime UTR | 0.0010 | 3 | 1.88 | 4.48E-01 |
| γ-glutamyltransferase | *GGT1* | Exon CADD | chr22:24603469 | - | C/T | 5-prime UTR | 0.0003 | 1 | 3.67 | 3.19E-01 |
| γ-glutamyltransferase | *GGT1* | Exon CADD | chr22:24607690 | rs61541228 | C/T | 5-prime UTR | 0.0151 | 44 | 4.64 | 9.27E-01 |
| γ-glutamyltransferase | *GGT1* | Exon CADD | chr22:24609020 | rs57719575 | G/C | 5-prime UTR | 0.0154 | 45 | 0.04 | 8.64E-01 |
| γ-glutamyltransferase | *GGT1* | Exon CADD | chr22:24609116 | - | G/A | 5-prime UTR | 0.0319 | 93 | 3.56 | 4.95E-01 |
| γ-glutamyltransferase | *GGT1* | Exon CADD | chr22:24609217 | rs549561372 | A/G | 5-prime UTR | 0.0003 | 1 | 0.50 | 1.08E-01 |
| γ-glutamyltransferase | *GGT1* | Exon CADD | chr22:24610277 | rs574639855 | T/C | 5-prime UTR | 0.0072 | 21 | 1.02 | 1.65E-01 |
| γ-glutamyltransferase | *GGT1* | Exon CADD | chr22:24610282 | - | G/A | 5-prime UTR | 0.0021 | 6 | 1.84 | 7.15E-01 |
| γ-glutamyltransferase | *GGT1* | Exon CADD | chr22:24610292 | rs541914297 | G/A | 5-prime UTR | 0.0185 | 54 | 1.36 | 8.51E-01 |
| γ-glutamyltransferase | *GGT1* | Exon CADD | chr22:24611165 | rs4049897 | A/G | Synonymous | 0.0141 | 41 | 0.02 | 4.98E-01 |
| γ-glutamyltransferase | *GGT1* | Exon CADD | chr22:24614791 | rs191957455 | C/T | Synonymous | 0.0075 | 22 | 11.72 | 2.01E-01 |
| γ-glutamyltransferase | *GGT1* | Exon CADD | chr22:24614792 | rs760767294 | G/A | Missense | 0.0038 | 11 | 23.80 | 4.38E-03 |
| γ-glutamyltransferase | *GGT1* | Exon CADD | chr22:24614888 | rs186765281 | A/G | Missense | 0.0202 | 59 | 8.80 | 2.98E-03 |
| γ-glutamyltransferase | *GGT1* | Exon CADD | chr22:24615095 | - | C/G | Missense | 0.0089 | 26 | 0.00 | 3.37E-01 |
| γ-glutamyltransferase | *GGT1* | Exon CADD | chr22:24615889 | rs2330848 | C/T | Non-coding transcript exon | 0.0034 | 10 | 0.19 | 2.94E-01 |
| γ-glutamyltransferase | *GGT1* | Exon CADD | chr22:24620331 | - | G/C | Missense | 0.0007 | 2 | 23.30 | 1.31E-01 |
| γ-glutamyltransferase | *GGT1* | Exon CADD | chr22:24620369 | rs200015790 | G/A | Missense | 0.0003 | 1 | 12.42 | 5.56E-01 |
| γ-glutamyltransferase | *GGT1* | Exon CADD | chr22:24621069 | - | C/T | Splice region | 0.0196 | 57 | 11.17 | 5.39E-01 |
| γ-glutamyltransferase | *GGT1* | Exon CADD | chr22:24627272 | rs531542863 | A/T | 5-prime UTR | 0.0017 | 5 | 0.78 | 3.20E-01 |
| γ-glutamyltransferase | *GGT1* | Exon CADD | chr22:24627279 | rs868669081 | G/A | Splice donor | 0.0082 | 24 | 4.97 | 3.42E-01 |
| γ-glutamyltransferase | *GGT1* | Exon CADD | chr22:24627365 | rs60762015 | C/T | 5-prime UTR | 0.0051 | 15 | 8.32 | 7.69E-01 |
| γ-glutamyltransferase | *GGT1* | Exon CADD | chr22:24627377 | rs554654292 | C/T | 5-prime UTR | 0.0251 | 73 | 16.32 | 1.28E-04 |
| γ-glutamyltransferase | *GGT1* | Exon CADD | chr22:24627439 | - | G/A | Missense | 0.0065 | 19 | 11.95 | 2.82E-01 |
| γ-glutamyltransferase | *GGT1* | Exon CADD | chr22:24627525 | rs71318991 | G/A | Missense | 0.0017 | 5 | 15.87 | 1.53E-01 |
| γ-glutamyltransferase | *GGT1* | Exon CADD | chr22:24627539 | rs201399186 | G/A | Synonymous | 0.0051 | 15 | 2.02 | 6.51E-01 |
| γ-glutamyltransferase | *GGT1* | Exon CADD | chr22:24627547 | rs781743521 | G/A | Missense | 0.0003 | 1 | 5.33 | 8.10E-03 |
| γ-glutamyltransferase | *GGT1* | Exon CADD | chr22:24627882 | rs200019915 | C/T | Synonymous | 0.0010 | 3 | 12.07 | 1.78E-01 |
| γ-glutamyltransferase | *GGT1* | Exon CADD | chr22:24628250 | rs372079440 | T/C | Synonymous | 0.0010 | 3 | 3.27 | 5.12E-01 |
| γ-glutamyltransferase | *GGT1* | Exon CADD | chr22:24628919 | rs149933133 | G/A | 3-prime UTR | 0.0096 | 28 | 1.12 | 4.41E-02 |

**Supplementary Table 3**

**Variants included in the burdens of the selected associations in the replication cohort.**

Alleles: reference/non-reference alleles; Consequence: the most severe consequence predicted by Ensembl VEP; MAF: minor allele frequency; AC: non-reference allele count; Weight: the weight assigned to the variant in the most significant test as per Supplementary Figure 8; NA indicates that no weight was applied.

| Trait | Gene | Analysis | Variant ID (GRCh38) | rsID | Alleles | Consequence | MAF | AC | Weight | single-point *P*-value |
|---|---|---|---|---|---|---|---|---|---|---|
| Triglycerides | *FAM189B* | LOFTEE LC | chr1:155250417 | rs749626426 | G/A | stop_gained | 0.0003 | 2 | NA | 5.35E-03 |
| Triglycerides | *FAM189B* | LOFTEE LC | chr1:155253873 | - | G/A | stop_gained | 0.0001 | 1 | NA | 5.88E-01 |
| Triglycerides | *APOC3* | Exon CADD+50 | chr11:116830638 | rs138326449 | G/A | splice_donor_variant | 0.0024 | 18 | 25.1 | 1.33E-04 |
| Triglycerides | *APOC3* | Exon CADD+50 | chr11:116829971 | rs191196015 | G/A | intron_variant | 0.0003 | 2 | 8.057 | 6.64E-01 |
| Triglycerides | *APOC3* | Exon CADD+50 | chr11:116829830 | . | C/G | 5_prime_UTR_variant | 0.0001 | 1 | 11.9 | 1.17E-02 |
| Triglycerides | *APOC3* | Exon CADD+50 | chr11:116831002 | rs527591419 | C/T | intron_variant | 0.0001 | 1 | 0.62 | 1.17E-02 |
| Triglycerides | *APOC3* | Exon CADD+50 | chr11:116830483 | rs748494867 | T/A | intron_variant | 0.0003 | 2 | 4.914 | 5.94E-02 |
| Triglycerides | *APOC3* | Exon CADD+50 | chr11:116830593 | rs779597455 | G/A | missense_variant | 0.0001 | 1 | 21.4 | 1.91E-01 |
| Triglycerides | *APOC3* | Exon CADD+50 | chr11:116830728 | rs184359086 | C/T | intron_variant | 0.0001 | 1 | 4.087 | 2.17E-01 |
| Triglycerides | *APOC3* | Exon CADD+50 | chr11:116831132 | . | C/T | intron_variant | 0.0003 | 2 | 0.361 | 3.99E-01 |
| Triglycerides | *APOC3* | Exon CADD+50 | chr11:116832848 | rs772522961 | C/A | missense_variant | 0.0001 | 1 | 1.016 | 8.23E-01 |
| Triglycerides | *APOC3* | Exon CADD+50 | chr11:116830844 | rs147210663 | G/A | missense_variant | 0.0005 | 4 | 23.6 | 5.16E-01 |
| Triglycerides | *APOC3* | Exon CADD+50 | chr11:116833039 | . | C/T | 3_prime_UTR_variant | 0.0001 | 1 | 10.18 | 5.00E-01 |
| Triglycerides | *APOC3* | Exon CADD+50 | chr11:116830637 | rs76353203 | C/T | stop_gained | 0.0007 | 5 | 32 | 2.42E-02 |
| Triglycerides | *APOC3* | Exon CADD+50 | chr11:116829929 | . | C/T | 5_prime_UTR_variant | 0.0001 | 1 | 12.84 | 5.76E-01 |
| Triglycerides | *APOC3* | Exon CADD+50 | chr11:116832764 | rs760769978 | G/A | splice_region_variant | 0.0001 | 1 | 3.119 | 7.19E-01 |
| Triglycerides | *APOC3* | Exon CADD+50 | chr11:116831090 | . | T/C | non_coding_transcript_exon_variant | 0.0005 | 4 | 2.332 | 7.33E-01 |
| Triglycerides | *APOC3* | Exon CADD+50 | chr11:116829772 | rs12721091 | G/A | 5_prime_UTR_variant | 0.0009 | 7 | 5.165 | 9.80E-01 |
| Triglycerides | *APOC3* | Exon CADD+50 | chr11:116830500 | rs5143 | G/A | 5_prime_UTR_variant | 0.0008 | 6 | 2.684 | 9.20E-01 |
| Triglycerides | *APOC3* | Exon CADD+50 | chr11:116832980 | rs577085953 | C/T | 3_prime_UTR_variant | 0.0001 | 1 | 8.906 | 1.56E-01 |
| Triglycerides | *APOC3* | Exon CADD+50 | chr11:116830843 | rs533891893 | C/T | synonymous_variant | 0.0001 | 1 | 6.244 | 1.32E-01 |
| Triglycerides | *APOC3* | Exon CADD+50 | chr11:116830620 | rs772815802 | C/T | missense_variant | 0.0001 | 1 | 10.97 | 7.77E-01 |
| Triglycerides | *APOC3* | Exon CADD+50 | chr11:116832729 | rs754046943 | A/T | intron_variant | 0.0001 | 1 | 1.848 | 6.88E-02 |
| Triglycerides | *APOC3* | Exon CADD+50 | chr11:116829771 | rs148295370 | C/T | 5_prime_UTR_variant | 0.0034 | 25 | 6.127 | 4.21E-01 |
| Triglycerides | *APOC3* | Exon CADD+50 | chr11:116831117 | rs1269330 | G/A | intron_variant | 0.0416 | 308 | 3.837 | 8.99E-01 |
| Triglycerides | *APOC3* | Exon CADD+50 | chr11:116833023 | rs187628630 | C/G | 3_prime_UTR_variant | 0.0071 | 53 | 7.365 | 8.98E-04 |
| Triglycerides | *APOC3* | Exon CADD+50 | chr11:116831133 | . | G/A | intron_variant | 0.0001 | 1 | 1.052 | 9.49E-01 |
| Bilirubin | *UGT1A9* | Exon+reg. EigenPhred | chr2:233672522 | rs151216459 | G/T | synonymous_variant | 0.0003 | 2 | 3.08850926 | 6.23E-01 |
| Bilirubin | *UGT1A9* | Exon+reg. EigenPhred | chr2:233712590 | . | A/G | intron_variant | 0.0003 | 2 | 2.341572053 | 6.00E-01 |
| Bilirubin | *UGT1A9* | Exon+reg. EigenPhred | chr2:233696085 | . | G/A | intron_variant | 0.0003 | 2 | 2.625818024 | 4.04E-01 |
| Bilirubin | *UGT1A9* | Exon+reg. EigenPhred | chr2:233773164 | rs768182403 | A/C | 3_prime_UTR_variant | 0.0001 | 1 | 3.873132917 | 7.25E-02 |
| Bilirubin | *UGT1A9* | Exon+reg. EigenPhred | chr2:233705432 | rs778048024 | A/G | intron_variant | 0.0001 | 1 | 0.075500585 | 6.31E-01 |
| Bilirubin | *UGT1A9* | Exon+reg. EigenPhred | chr2:233704927 | . | C/T | intron_variant | 0.0001 | 1 | 3.171551397 | 2.15E-01 |
| Bilirubin | *UGT1A9* | Exon+reg. EigenPhred | chr2:233744758 | . | T/C | intron_variant | 0.0001 | 1 | 6.958886244 | 4.76E-03 |
| Bilirubin | *UGT1A9* | Exon+reg. EigenPhred | chr2:233702413 | rs182081835 | T/G | intron_variant | 0.0051 | 38 | 0.09719573 | 1.08E-01 |
| Bilirubin | *UGT1A9* | Exon+reg. EigenPhred | chr2:233676786 | rs74435166 | C/G | intron_variant | 0.0195 | 145 | 4.117716181 | 8.47E-02 |
| Bilirubin | *UGT1A9* | Exon+reg. EigenPhred | chr2:233766051 | . | C/G | intron_variant | 0.0001 | 1 | 2.275392907 | 5.40E-01 |
| Bilirubin | *UGT1A9* | Exon+reg. EigenPhred | chr2:233702319 | . | C/T | intron_variant | 0.0001 | 1 | 1.913812296 | 9.46E-01 |
| Bilirubin | *UGT1A9* | Exon+reg. EigenPhred | chr2:233755491 | rs112011393 | T/C | non_coding_transcript_exon_variant | 0.0004 | 3 | 3.017253775 | 1.07E-01 |
| Bilirubin | *UGT1A9* | Exon+reg. EigenPhred | chr2:233766771 | rs71528513 | C/T | intron_variant | 0.0081 | 60 | 4.686024065 | 1.09E-01 |
| Bilirubin | *UGT1A9* | Exon+reg. EigenPhred | chr2:233676802 | . | G/A | intron_variant | 0.0001 | 1 | 1.720037511 | 1.90E-02 |
| Bilirubin | *UGT1A9* | Exon+reg. EigenPhred | chr2:233766201 | rs544565545 | G/T | intron_variant | 0.0003 | 2 | 0.978756042 | 5.31E-01 |
| Bilirubin | *UGT1A9* | Exon+reg. EigenPhred | chr2:233712668 | . | G/A | intron_variant | 0.0001 | 1 | 6.19701633 | 9.02E-01 |
| Bilirubin | *UGT1A9* | Exon+reg. EigenPhred | chr2:233712705 | . | T/C | intron_variant | 0.0001 | 1 | 4.697412975 | 2.48E-01 |
| Bilirubin | *UGT1A9* | Exon+reg. EigenPhred | chr2:233674892 | rs45488098 | T/C | intron_variant | 0.0052 | 39 | 3.188918433 | 1.84E-01 |
| Bilirubin | *UGT1A9* | Exon+reg. EigenPhred | chr2:233679436 | rs752875090 | C/G | intron_variant | 0.0001 | 1 | 0.061626737 | 1.90E-01 |
| Bilirubin | *UGT1A9* | Exon+reg. EigenPhred | chr2:233761138 | . | A/G | missense_variant | 0.0001 | 1 | 0.002796919 | 8.49E-01 |
| Bilirubin | *UGT1A9* | Exon+reg. EigenPhred | chr2:233672196 | rs143487779 | C/T | missense_variant | 0.0016 | 12 | 0.626982503 | 5.59E-01 |
| Bilirubin | *UGT1A9* | Exon+reg. EigenPhred | chr2:233692704 | . | C/G | intron_variant | 0.0001 | 1 | 8.679314838 | 7.46E-01 |
| Bilirubin | *UGT1A9* | Exon+reg. EigenPhred | chr2:233766325 | rs574966930 | G/A | intron_variant | 0.0004 | 3 | 0.188510611 | 9.81E-01 |
| Bilirubin | *UGT1A9* | Exon+reg. EigenPhred | chr2:233695529 | rs774353867 | T/C | intron_variant | 0.0001 | 1 | 7.596230891 | 6.53E-01 |
| Bilirubin | *UGT1A9* | Exon+reg. EigenPhred | chr2:233755421 | rs774443994 | C/T | non_coding_transcript_exon_variant | 0.0009 | 7 | 0.926052068 | 5.27E-02 |
| Bilirubin | *UGT1A9* | Exon+reg. EigenPhred | chr2:233679517 | . | C/T | intron_variant | 0.0001 | 1 | 1.350445925 | 2.51E-01 |
| Bilirubin | *UGT1A9* | Exon+reg. EigenPhred | chr2:233672237 | . | A/G | synonymous_variant | 0.0001 | 1 | 1.807427221 | 6.13E-01 |
| Bilirubin | *UGT1A9* | Exon+reg. EigenPhred | chr2:233755420 | rs187061066 | G/A | non_coding_transcript_exon_variant | 0.0003 | 2 | 2.241848571 | 9.43E-01 |
| Bilirubin | *UGT1A9* | Exon+reg. EigenPhred | chr2:233767212 | rs377453564 | G/A | intron_variant | 0.0004 | 3 | 6.297725591 | 5.95E-01 |
| Bilirubin | *UGT1A9* | Exon+reg. EigenPhred | chr2:233697105 | . | G/T | intron_variant | 0.0001 | 1 | 1.307693515 | 6.99E-01 |
| Bilirubin | *UGT1A9* | Exon+reg. EigenPhred | chr2:233709696 | rs554703505 | T/C | intron_variant | 0.0001 | 1 | 0.379387193 | 4.53E-01 |
| Bilirubin | *UGT1A9* | Exon+reg. EigenPhred | chr2:233692105 | . | A/G | 3_prime_UTR_variant | 0.0001 | 1 | 5.083110604 | 3.15E-01 |
| Bilirubin | *UGT1A9* | Exon+reg. EigenPhred | chr2:233675321 | rs746332886 | T/C | intron_variant | 0.0001 | 1 | 1.392263388 | 1.42E-01 |
| Bilirubin | *UGT1A9* | Exon+reg. EigenPhred | chr2:233675068 | rs549929668 | T/C | intron_variant | 0.0001 | 1 | 2.431502275 | 7.71E-01 |
| Bilirubin | *UGT1A9* | Exon+reg. EigenPhred | chr2:233711177 | rs141849516 | G/A | intron_variant | 0.0005 | 4 | 0.284941132 | 6.84E-01 |
| Bilirubin | *UGT1A9* | Exon+reg. EigenPhred | chr2:233705460 | . | C/T | intron_variant | 0.0001 | 1 | 3.97764383 | 5.78E-01 |
| Bilirubin | *UGT1A9* | Exon+reg. EigenPhred | chr2:233702494 | rs374252732 | C/T | intron_variant | 0.0013 | 10 | 0.000817043 | 4.85E-02 |
| Bilirubin | *UGT1A9* | Exon+reg. EigenPhred | chr2:233705319 | rs112242313 | A/G | intron_variant | 0.0008 | 6 | 0.570785304 | 3.26E-01 |
| Bilirubin | *UGT1A9* | Exon+reg. EigenPhred | chr2:233767971 | rs71537802 | A/C | intron_variant | 0.0008 | 6 | 6.391019145 | 2.78E-02 |

| | | | | | | | | | | |
|---|---|---|---|---|---|---|---|---|---|---|
| Bilirubin | *UGT1A9* | Exon+reg. EigenPhred | chr2:233744794 | rs145652010 | G/C | intron_variant | 0.0003 | 2 | 3.80087609 | 7.37E-01 |
| Bilirubin | *UGT1A9* | Exon+reg. EigenPhred | chr2:233697767 | . | G/T | intron_variant | 0.0001 | 1 | 3.219785857 | 7.94E-02 |
| Bilirubin | *UGT1A9* | Exon+reg. EigenPhred | chr2:233744646 | rs28899468 | T/C | intron_variant | 0.0313 | 233 | 2.743433159 | 1.47E-16 |
| Bilirubin | *UGT1A9* | Exon+reg. EigenPhred | chr2:233767175 | . | T/G | intron_variant | 0.0001 | 1 | 9.615572359 | 3.17E-01 |
| Bilirubin | *UGT1A9* | Exon+reg. EigenPhred | chr2:233760481 | rs762109713 | C/T | missense_variant | 0.0003 | 2 | 2.017335156 | 8.14E-01 |
| Bilirubin | *UGT1A9* | Exon+reg. EigenPhred | chr2:233714126 | . | C/T | intron_variant | 0.0001 | 1 | 3.610665434 | 1.24E-01 |
| Bilirubin | *UGT1A9* | Exon+reg. EigenPhred | chr2:233709733 | rs190992275 | T/C | intron_variant | 0.0058 | 43 | 2.146237167 | 2.42E-01 |
| Bilirubin | *UGT1A9* | Exon+reg. EigenPhred | chr2:233674107 | . | A/G | intron_variant | 0.0001 | 1 | 0.15137646 | 6.70E-01 |
| Bilirubin | *UGT1A9* | Exon+reg. EigenPhred | chr2:233702289 | . | A/G | intron_variant | 0.0001 | 1 | 0.557858968 | 8.59E-01 |
| Bilirubin | *UGT1A9* | Exon+reg. EigenPhred | chr2:233702170 | . | C/T | intron_variant | 0.0003 | 2 | 1.241751991 | 6.86E-02 |
| Bilirubin | *UGT1A9* | Exon+reg. EigenPhred | chr2:233744892 | rs28900378 | C/T | intron_variant | 0.0009 | 7 | 0.045840174 | 8.80E-01 |
| Bilirubin | *UGT1A9* | Exon+reg. EigenPhred | chr2:233772475 | rs190427553 | C/A | synonymous_variant | 0.0003 | 2 | 13.95092946 | 4.66E-02 |
| Bilirubin | *UGT1A9* | Exon+reg. EigenPhred | chr2:233745061 | rs115414076 | T/C | intron_variant | 0.0274 | 204 | 1.586437596 | 2.35E-16 |
| Bilirubin | *UGT1A9* | Exon+reg. EigenPhred | chr2:233674127 | . | T/C | intron_variant | 0.0003 | 2 | 5.341662693 | 7.82E-01 |
| Bilirubin | *UGT1A9* | Exon+reg. EigenPhred | chr2:233744492 | . | G/A | intron_variant | 0.0001 | 1 | 1.148745586 | 6.72E-01 |
| Bilirubin | *UGT1A9* | Exon+reg. EigenPhred | chr2:233691801 | . | C/G | 5_prime_UTR_variant | 0.0001 | 1 | 4.740433552 | 9.87E-02 |
| Bilirubin | *UGT1A9* | Exon+reg. EigenPhred | chr2:233676925 | rs28970011 | G/A | intron_variant | 0.0003 | 2 | 4.076224978 | 5.31E-01 |
| Bilirubin | *UGT1A9* | Exon+reg. EigenPhred | chr2:233674300 | rs74471804 | C/T | intron_variant | 0.0001 | 1 | 0.282986669 | 7.04E-01 |
| Bilirubin | *UGT1A9* | Exon+reg. EigenPhred | chr2:233671853 | . | G/T | 5_prime_UTR_variant | 0.0001 | 1 | 7.133029349 | 1.19E-01 |
| Bilirubin | *UGT1A9* | Exon+reg. EigenPhred | chr2:233766252 | . | C/T | intron_variant | 0.0003 | 2 | 0.095726348 | 4.10E-01 |
| Bilirubin | *UGT1A9* | Exon+reg. EigenPhred | chr2:233714147 | rs548246419 | C/T | intron_variant | 0.0001 | 1 | 4.993753562 | 1.56E-01 |
| Bilirubin | *UGT1A9* | Exon+reg. EigenPhred | chr2:233702197 | rs187320668 | A/T | intron_variant | 0.0004 | 3 | 2.901113267 | 5.06E-01 |
| Bilirubin | *UGT1A9* | Exon+reg. EigenPhred | chr2:233702653 | rs746758622 | G/A | intron_variant | 0.0009 | 7 | 0.118609082 | 3.57E-01 |
| Bilirubin | *UGT1A9* | Exon+reg. EigenPhred | chr2:233672434 | rs151238339 | T/C | missense_variant | 0.0004 | 3 | 0.117675386 | 4.59E-01 |
| Bilirubin | *UGT1A9* | Exon+reg. EigenPhred | chr2:233672460 | rs141811044 | T/C | missense_variant | 0.0009 | 7 | 0.042740382 | 1.20E-03 |
| Bilirubin | *UGT1A9* | Exon+reg. EigenPhred | chr2:233767864 | rs748591879 | A/G | missense_variant | 0.0001 | 1 | 11.5611763 | 5.77E-02 |
| Bilirubin | *UGT1A9* | Exon+reg. EigenPhred | chr2:233766381 | . | T/G | intron_variant | 0.0003 | 2 | 0.193675167 | 1.91E-01 |
| Bilirubin | *UGT1A9* | Exon+reg. EigenPhred | chr2:233696549 | . | T/C | intron_variant | 0.0003 | 2 | 2.956091704 | 6.80E-01 |
| Bilirubin | *UGT1A9* | Exon+reg. EigenPhred | chr2:233705352 | . | A/G | intron_variant | 0.0001 | 1 | 0.956858866 | 2.46E-01 |
| Bilirubin | *UGT1A9* | Exon+reg. EigenPhred | chr2:233762032 | rs34562642 | T/A | intron_variant | 0.0008 | 6 | 0.093750497 | 3.37E-01 |
| Bilirubin | *UGT1A9* | Exon+reg. EigenPhred | chr2:233702394 | rs767929704 | T/C | intron_variant | 0.0001 | 1 | 0.739704076 | 3.29E-01 |
| Bilirubin | *UGT1A9* | Exon+reg. EigenPhred | chr2:233772427 | rs114123636 | C/T | synonymous_variant | 0.0001 | 1 | 12.351444 | 6.14E-01 |
| Bilirubin | *UGT1A9* | Exon+reg. EigenPhred | chr2:233705496 | rs190154138 | C/A | intron_variant | 0.0047 | 35 | 0.230173477 | 9.16E-02 |
| Bilirubin | *UGT1A9* | Exon+reg. EigenPhred | chr2:233766327 | rs373926451 | G/A | intron_variant | 0.0001 | 1 | 0.155638982 | 8.12E-01 |
| Bilirubin | *UGT1A9* | Exon+reg. EigenPhred | chr2:233772335 | rs115410088 | T/C | missense_variant | 0.0001 | 1 | 13.03260395 | 3.45E-01 |
| Bilirubin | *UGT1A9* | Exon+reg. EigenPhred | chr2:233755844 | . | G/T | non_coding_transcript_exon_variant | 0.0001 | 1 | 0.174782446 | 7.40E-01 |
| Bilirubin | *UGT1A9* | Exon+reg. EigenPhred | chr2:233695470 | rs763759412 | T/C | intron_variant | 0.0001 | 1 | 4.250301017 | 3.97E-01 |
| Bilirubin | *UGT1A9* | Exon+reg. EigenPhred | chr2:233695283 | rs546580338 | T/A | intron_variant | 0.0004 | 3 | 0.074573367 | 7.23E-03 |
| Bilirubin | *UGT1A9* | Exon+reg. EigenPhred | chr2:233698470 | . | T/A | intron_variant | 0.0003 | 2 | 3.245730499 | 8.78E-01 |
| Bilirubin | *UGT1A9* | Exon+reg. EigenPhred | chr2:233695902 | rs184289698 | G/T | intron_variant | 0.0001 | 1 | 0.169404793 | 7.44E-01 |
| Bilirubin | *UGT1A9* | Exon+reg. EigenPhred | chr2:233705656 | . | A/G | intron_variant | 0.0001 | 1 | 1.325594927 | 3.71E-01 |
| Bilirubin | *UGT1A9* | Exon+reg. EigenPhred | chr2:233672660 | rs66915469 | T/G | stop_gained | 0.0001 | 1 | 1.437057991 | 8.84E-01 |
| Bilirubin | *UGT1A9* | Exon+reg. EigenPhred | chr2:233676622 | rs77410236 | C/T | intron_variant | 0.0001 | 1 | 0.133650653 | 9.62E-01 |
| Bilirubin | *UGT1A9* | Exon+reg. EigenPhred | chr2:233680786 | rs542991668 | G/A | intron_variant | 0.0001 | 1 | 4.764191779 | 6.68E-01 |
| Bilirubin | *UGT1A9* | Exon+reg. EigenPhred | chr2:233705548 | . | G/C | intron_variant | 0.0001 | 1 | 1.584557602 | 6.28E-01 |
| Bilirubin | *UGT1A9* | Exon+reg. EigenPhred | chr2:233761493 | rs34652311 | C/T | intron_variant | 0.0009 | 7 | 8.506433637 | 8.80E-01 |
| Bilirubin | *UGT1A9* | Exon+reg. EigenPhred | chr2:233761235 | rs35041092 | C/T | intron_variant | 0.0012 | 9 | 10.26524394 | 1.93E-01 |
| Bilirubin | *UGT1A9* | Exon+reg. EigenPhred | chr2:233745103 | . | T/C | intron_variant | 0.0001 | 1 | 0.483756128 | 2.15E-01 |
| Bilirubin | *UGT1A9* | Exon+reg. EigenPhred | chr2:233712790 | rs28898600 | T/G | intron_variant | 0.0009 | 7 | 0.725994967 | 8.80E-01 |
| Bilirubin | *UGT1A9* | Exon+reg. EigenPhred | chr2:233692572 | rs757852696 | G/A | intron_variant | 0.0007 | 5 | 4.575464286 | 6.13E-01 |
| Bilirubin | *UGT1A9* | Exon+reg. EigenPhred | chr2:233736818 | rs761920295 | G/A | intron_variant | 0.0001 | 1 | 0.118983171 | 5.22E-01 |
| Bilirubin | *UGT1A9* | Exon+reg. EigenPhred | chr2:233697448 | rs780047517 | C/G | intron_variant | 0.0003 | 2 | 0.450767401 | 4.72E-01 |
| Bilirubin | *UGT1A9* | Exon+reg. EigenPhred | chr2:233697097 | . | T/C | intron_variant | 0.0001 | 1 | 2.74864941 | 5.56E-01 |
| Bilirubin | *UGT1A9* | Exon+reg. EigenPhred | chr2:233672032 | rs72551330 | T/C | missense_variant | 0.0164 | 122 | 0.714044824 | 3.94E-01 |
| Bilirubin | *UGT1A9* | Exon+reg. EigenPhred | chr2:233712842 | rs11689628 | C/T | intron_variant | 0.0267 | 199 | 1.180305293 | 1.67E-16 |
| Bilirubin | *UGT1A9* | Exon+reg. EigenPhred | chr2:233695871 | rs182203911 | C/T | intron_variant | 0.0024 | 18 | 1.458050903 | 1.89E-01 |
| Bilirubin | *UGT1A9* | Exon+reg. EigenPhred | chr2:233772309 | rs114982090 | C/T | missense_variant | 0.0001 | 1 | 14.55166778 | 3.02E-01 |
| Bilirubin | *UGT1A9* | Exon+reg. EigenPhred | chr2:233711259 | . | C/G | intron_variant | 0.0001 | 1 | 1.441782797 | 9.80E-01 |
| Bilirubin | *UGT1A9* | Exon+reg. EigenPhred | chr2:233680543 | rs184094797 | C/G | intron_variant | 0.0003 | 2 | 2.111036501 | 3.54E-01 |
| Bilirubin | *UGT1A9* | Exon+reg. EigenPhred | chr2:233672430 | rs766064889 | G/A | missense_variant | 0.0001 | 1 | 10.17078153 | 6.17E-01 |
| Bilirubin | *UGT1A9* | Exon+reg. EigenPhred | chr2:233676898 | . | G/A | intron_variant | 0.0001 | 1 | 0.119606723 | 6.54E-01 |
| Bilirubin | *UGT1A9* | Exon+reg. EigenPhred | chr2:233755391 | rs115647781 | C/G | non_coding_transcript_exon_variant | 0.0113 | 84 | 5.107243052 | 1.85E-06 |
| Bilirubin | *UGT1A9* | Exon+reg. EigenPhred | chr2:233697701 | rs765701149 | G/T | intron_variant | 0.0001 | 1 | 0.66470667 | 7.26E-01 |
| Bilirubin | *UGT1A9* | Exon+reg. EigenPhred | chr2:233755509 | rs143373661 | C/G | non_coding_transcript_exon_variant | 0.0452 | 337 | 6.018128149 | 3.14E-18 |
| Bilirubin | *UGT1A9* | Exon+reg. EigenPhred | chr2:233765924 | . | G/A | intron_variant | 0.0001 | 1 | 0.324004168 | 1.76E-01 |
| Bilirubin | *UGT1A9* | Exon+reg. EigenPhred | chr2:233680663 | . | G/C | intron_variant | 0.0003 | 2 | 3.036540856 | 7.09E-01 |
| Bilirubin | *UGT1A9* | Exon+reg. EigenPhred | chr2:233676479 | rs34707303 | C/T | intron_variant | 0.0121 | 90 | 1.131186229 | 1.59E-02 |
| Bilirubin | *UGT1A9* | Exon+reg. EigenPhred | chr2:233762169 | . | G/A | intron_variant | 0.0004 | 3 | 0.494094236 | 3.90E-02 |
| Bilirubin | *UGT1A9* | Exon+reg. EigenPhred | chr2:233675200 | rs12615708 | G/A | intron_variant | 0.0236 | 176 | 0.158764509 | 1.91E-04 |
| Bilirubin | *UGT1A9* | Exon+reg. EigenPhred | chr2:233679365 | rs754142123 | T/C | intron_variant | 0.0003 | 2 | 0.655581927 | 4.47E-01 |
| Bilirubin | *UGT1A9* | Exon+reg. EigenPhred | chr2:233692098 | . | G/C | 3_prime_UTR_variant | 0.0001 | 1 | 0.227641092 | 9.66E-01 |
| Bilirubin | *UGT1A9* | Exon+reg. EigenPhred | chr2:233767880 | rs144978321 | C/T | missense_variant | 0.0001 | 1 | 7.956440915 | 2.89E-01 |
| Bilirubin | *UGT1A9* | Exon+reg. EigenPhred | chr2:233709576 | rs180747204 | A/C | intron_variant | 0.0009 | 7 | 1.69090005 | 3.46E-01 |
| Bilirubin | *UGT1A9* | Exon+reg. EigenPhred | chr2:233676481 | . | C/A | intron_variant | 0.0003 | 2 | 0.989280569 | 5.95E-01 |
| Bilirubin | *UGT1A9* | Exon+reg. EigenPhred | chr2:233676666 | rs768806878 | G/A | intron_variant | 0.0001 | 1 | 1.565249385 | 6.33E-01 |
| Bilirubin | *UGT1A9* | Exon+reg. EigenPhred | chr2:233692029 | . | G/C | 3_prime_UTR_variant | 0.0003 | 2 | 7.071421945 | 3.74E-01 |
| Bilirubin | *UGT1A9* | Exon+reg. EigenPhred | chr2:233711354 | . | C/G | intron_variant | 0.0004 | 3 | 0.411471525 | 4.61E-02 |
| Bilirubin | *UGT1A9* | Exon+reg. EigenPhred | chr2:233674004 | rs17862858 | A/G | intron_variant | 0.0193 | 144 | 2.195623312 | 1.17E-01 |
| Bilirubin | *UGT1A9* | Exon+reg. EigenPhred | chr2:233672780 | rs767375950 | G/A | synonymous_variant | 0.0001 | 1 | 3.966251492 | 3.21E-01 |
| Bilirubin | *UGT1A9* | Exon+reg. EigenPhred | chr2:233697391 | rs74792728 | T/A | intron_variant | 0.0001 | 1 | 3.730686514 | 4.19E-01 |
| Bilirubin | *UGT1A9* | Exon+reg. EigenPhred | chr2:233697371 | . | T/C | intron_variant | 0.0004 | 3 | 2.570302964 | 3.54E-01 |

| | | | | | | | | | | |
|---|---|---|---|---|---|---|---|---|---|---|
| Bilirubin | *UGT1A9* | Exon+reg. EigenPhred | chr2:233675244 | . | G/A | intron_variant | 0.0001 | 1 | 3.277401502 | 3.18E-01 |
| Bilirubin | *UGT1A9* | Exon+reg. EigenPhred | chr2:233675250 | rs757802955 | C/T | intron_variant | 0.0003 | 2 | 2.68498789 | 4.20E-01 |
| Bilirubin | *UGT1A9* | Exon+reg. EigenPhred | chr2:233747264 | . | A/G | non_coding_transcript_exon_variant | 0.0003 | 2 | 1.470520006 | 6.80E-01 |
| Bilirubin | *UGT1A9* | Exon+reg. EigenPhred | chr2:233767183 | rs34650714 | C/T | intron_variant | 0.0003 | 2 | 2.03171021 | 7.37E-01 |
| Bilirubin | *UGT1A9* | Exon+reg. EigenPhred | chr2:233772677 | . | A/C | 3_prime_UTR_variant | 0.0001 | 1 | 3.512604341 | 6.17E-01 |
| Bilirubin | *UGT1A9* | Exon+reg. EigenPhred | chr2:233762191 | . | G/A | intron_variant | 0.0001 | 1 | 2.908139218 | 5.67E-01 |
| Bilirubin | *UGT1A9* | Exon+reg. EigenPhred | chr2:233773055 | . | G/A | 3_prime_UTR_variant | 0.0003 | 2 | 2.900687338 | 3.00E-01 |
| Bilirubin | *UGT1A9* | Exon+reg. EigenPhred | chr2:233675421 | rs769866030 | T/C | intron_variant | 0.0003 | 2 | 1.134318501 | 1.32E-01 |
| Bilirubin | *UGT1A9* | Exon+reg. EigenPhred | chr2:233705012 | rs544165304 | T/A | intron_variant | 0.0016 | 12 | 1.207607357 | 1.33E-01 |
| Bilirubin | *UGT1A9* | Exon+reg. EigenPhred | chr2:233737048 | rs764914717 | T/C | intron_variant | 0.0004 | 3 | 1.499079268 | 1.43E-01 |
| Bilirubin | *UGT1A9* | Exon+reg. EigenPhred | chr2:233755783 | rs142854795 | C/G | non_coding_transcript_exon_variant | 0.0177 | 132 | 2.438483553 | 2.69E-02 |
| Bilirubin | *UGT1A9* | Exon+reg. EigenPhred | chr2:233698474 | rs17868329 | A/G | intron_variant | 0.0176 | 131 | 1.927041319 | 2.83E-05 |
| Bilirubin | *UGT1A9* | Exon+reg. EigenPhred | chr2:233674049 | rs753425003 | G/A | intron_variant | 0.0001 | 1 | 0.480806467 | 8.41E-01 |
| Bilirubin | *UGT1A9* | Exon+reg. EigenPhred | chr2:233692808 | . | G/A | intron_variant | 0.0001 | 1 | 7.522253988 | 4.32E-01 |
| Bilirubin | *UGT1A9* | Exon+reg. EigenPhred | chr2:233772241 | rs200350292 | C/T | intron_variant | 0.0001 | 1 | 6.566625608 | 9.57E-01 |
| Bilirubin | *UGT1A9* | Exon+reg. EigenPhred | chr2:233672783 | . | G/T | missense_variant | 0.0001 | 1 | 5.11354004 | 5.90E-01 |
| Bilirubin | *UGT1A9* | Exon+reg. EigenPhred | chr2:233679502 | rs187149830 | G/A | intron_variant | 0.0003 | 2 | 0.166179154 | 1.90E-02 |
| Bilirubin | *UGT1A9* | Exon+reg. EigenPhred | chr2:233712808 | . | G/A | intron_variant | 0.0001 | 1 | 2.0844113 | 7.35E-02 |
| Bilirubin | *UGT1A9* | Exon+reg. EigenPhred | chr2:233701753 | rs540599933 | A/G | intron_variant | 0.0001 | 1 | 0.439362184 | 3.56E-01 |
| Bilirubin | *UGT1A9* | Exon+reg. EigenPhred | chr2:233695767 | . | G/A | intron_variant | 0.0001 | 1 | 1.156483836 | 5.37E-01 |
| Bilirubin | *UGT1A9* | Exon+reg. EigenPhred | chr2:233760581 | rs138183896 | T/C | synonymous_variant | 0.0003 | 2 | 6.459431101 | 2.98E-01 |
| Bilirubin | *UGT1A9* | Exon+reg. EigenPhred | chr2:233758543 | . | T/C | intron_variant | 0.0001 | 1 | 3.56313082 | 5.52E-01 |
| Bilirubin | *UGT1A9* | Exon+reg. EigenPhred | chr2:233772729 | . | G/A | 3_prime_UTR_variant | 0.0003 | 2 | 1.014821774 | 5.62E-01 |
| Bilirubin | *UGT1A9* | Exon+reg. EigenPhred | chr2:233711204 | rs760514189 | C/T | intron_variant | 0.0008 | 6 | 0.106503245 | 7.78E-01 |
| Bilirubin | *UGT1A9* | Exon+reg. EigenPhred | chr2:233711088 | rs145698870 | A/G | intron_variant | 0.0003 | 2 | 0.288440637 | 7.37E-01 |
| Bilirubin | *UGT1A9* | Exon+reg. EigenPhred | chr2:233702369 | rs116955810 | G/A | intron_variant | 0.0079 | 59 | 0.098621478 | 1.55E-01 |
| Bilirubin | *UGT1A9* | Exon+reg. EigenPhred | chr2:233672109 | rs199829358 | A/C | missense_variant | 0.0003 | 2 | 0.005884521 | 9.53E-01 |
| Bilirubin | *UGT1A9* | Exon+reg. EigenPhred | chr2:233772705 | rs769331263 | C/G | 3_prime_UTR_variant | 0.0001 | 1 | 0.01666785 | 1.50E-01 |
| Bilirubin | *UGT1A9* | Exon+reg. EigenPhred | chr2:233680491 | rs142850766 | A/G | intron_variant | 0.0055 | 41 | 4.638285047 | 2.98E-01 |
| Bilirubin | *UGT1A9* | Exon+reg. EigenPhred | chr2:233712649 | . | G/A | intron_variant | 0.0001 | 1 | 5.128242272 | 4.29E-01 |
| Bilirubin | *UGT1A9* | Exon+reg. EigenPhred | chr2:233768257 | rs139698110 | T/C | synonymous_variant | 0.002 | 15 | 17.80625631 | 4.92E-01 |
| Bilirubin | *UGT1A9* | Exon+reg. EigenPhred | chr2:233766514 | . | T/C | intron_variant | 0.0001 | 1 | 1.82828071 | 7.12E-01 |
| Bilirubin | *UGT1A9* | Exon+reg. EigenPhred | chr2:233714390 | . | A/G | intron_variant | 0.0001 | 1 | 0.669608104 | 9.79E-01 |
| Bilirubin | *UGT1A9* | Exon+reg. EigenPhred | chr2:233772452 | rs199723856 | G/A | missense_variant | 0.0003 | 2 | 3.763554513 | 3.80E-01 |
| Bilirubin | *UGT1A9* | Exon+reg. EigenPhred | chr2:233772635 | . | A/G | 3_prime_UTR_variant | 0.0001 | 1 | 12.96051955 | 3.76E-01 |
| Bilirubin | *UGT1A9* | Exon+reg. EigenPhred | chr2:233767927 | rs267599273 | G/A | missense_variant | 0.0003 | 2 | 15.99439333 | 5.05E-02 |
| Bilirubin | *UGT1A9* | Exon+reg. EigenPhred | chr2:233709577 | rs186174292 | A/T | intron_variant | 0.0009 | 7 | 2.131967582 | 3.46E-01 |
| Bilirubin | *UGT1A9* | Exon+reg. EigenPhred | chr2:233695502 | . | G/T | intron_variant | 0.0001 | 1 | 7.393734817 | 5.84E-01 |
| Bilirubin | *UGT1A9* | Exon+reg. EigenPhred | chr2:233755384 | . | G/A | non_coding_transcript_exon_variant | 0.0001 | 1 | 15.81309389 | 2.45E-01 |
| Bilirubin | *UGT1A9* | Exon+reg. EigenPhred | chr2:233697058 | . | G/C | intron_variant | 0.0003 | 2 | 0.291112527 | 6.62E-01 |
| Bilirubin | *UGT1A9* | Exon+reg. EigenPhred | chr2:233705445 | rs79916929 | G/A | intron_variant | 0.0079 | 59 | 3.072391633 | 2.16E-01 |
| Bilirubin | *UGT1A9* | Exon+reg. EigenPhred | chr2:233696249 | . | A/C | intron_variant | 0.0001 | 1 | 1.282636035 | 3.96E-01 |
| Bilirubin | *UGT1A9* | Exon+reg. EigenPhred | chr2:233760498 | rs4148323 | G/A | missense_variant | 0.0009 | 7 | 3.764965783 | 6.80E-01 |
| Bilirubin | *UGT1A9* | Exon+reg. EigenPhred | chr2:233711341 | . | T/C | intron_variant | 0.0001 | 1 | 0.565321459 | 8.44E-01 |
| Bilirubin | *UGT1A9* | Exon+reg. EigenPhred | chr2:233705516 | rs769825689 | G/T | intron_variant | 0.0003 | 2 | 0.136323558 | 3.14E-01 |
| Bilirubin | *UGT1A9* | Exon+reg. EigenPhred | chr2:233705198 | . | T/C | intron_variant | 0.0001 | 1 | 1.662905664 | 6.19E-01 |
| Bilirubin | *UGT1A9* | Exon+reg. EigenPhred | chr2:233672286 | . | G/T | missense_variant | 0.0001 | 1 | 4.128139008 | 6.80E-02 |
| Bilirubin | *UGT1A9* | Exon+reg. EigenPhred | chr2:233674064 | . | A/C | intron_variant | 0.0001 | 1 | 2.687690534 | 2.18E-01 |
| Bilirubin | *UGT1A9* | Exon+reg. EigenPhred | chr2:233672273 | rs760653064 | C/T | synonymous_variant | 0.0003 | 2 | 3.161601036 | 9.31E-01 |
| Bilirubin | *UGT1A9* | Exon+reg. EigenPhred | chr2:233675462 | rs79913681 | C/G | intron_variant | 0.0189 | 141 | 0.464236472 | 6.42E-04 |
| Bilirubin | *UGT1A9* | Exon+reg. EigenPhred | chr2:233712259 | rs763507472 | G/A | intron_variant | 0.0007 | 5 | 5.209178769 | 2.32E-01 |
| Bilirubin | *UGT1A9* | Exon+reg. EigenPhred | chr2:233760693 | . | C/T | missense_variant | 0.0001 | 1 | 2.193688161 | 6.18E-01 |
| Bilirubin | *UGT1A9* | Exon+reg. EigenPhred | chr2:233760961 | rs35003977 | T/G | missense_variant | 0.0003 | 2 | 0.016264582 | 9.93E-01 |
| Bilirubin | *UGT1A9* | Exon+reg. EigenPhred | chr2:233761978 | . | G/A | intron_variant | 0.0001 | 1 | 0.809806218 | 2.57E-01 |
| Bilirubin | *UGT1A9* | Exon+reg. EigenPhred | chr2:233747270 | . | T/C | non_coding_transcript_exon_variant | 0.0003 | 2 | 1.614102142 | 6.80E-01 |
| Bilirubin | *UGT1A9* | Exon+reg. EigenPhred | chr2:233772828 | rs762669023 | C/T | 3_prime_UTR_variant | 0.0001 | 1 | 8.947758237 | 6.17E-01 |
| Bilirubin | *UGT1A9* | Exon+reg. EigenPhred | chr2:233692449 | rs45568235 | C/T | intron_variant | 0.0001 | 1 | 4.004665916 | 1.95E-01 |
| Bilirubin | *UGT1A9* | Exon+reg. EigenPhred | chr2:233672159 | rs41264153 | G/A | synonymous_variant | 0.0095 | 71 | 12.98082496 | 4.14E-02 |
| Bilirubin | *UGT1A9* | Exon+reg. EigenPhred | chr2:233744652 | rs754395729 | A/G | intron_variant | 0.0007 | 5 | 0.588570354 | 6.13E-01 |
| Bilirubin | *UGT1A9* | Exon+reg. EigenPhred | chr2:233761536 | . | A/C | intron_variant | 0.0001 | 1 | 4.794947729 | 6.02E-01 |
| Bilirubin | *UGT1A9* | Exon+reg. EigenPhred | chr2:233672501 | rs373933295 | T/C | synonymous_variant | 0.0001 | 1 | 8.82043369 | 6.13E-01 |
| Bilirubin | *UGT1A9* | Exon+reg. EigenPhred | chr2:233755371 | . | G/A | non_coding_transcript_exon_variant | 0.0001 | 1 | 1.145497592 | 4.79E-01 |
| Bilirubin | *UGT1A9* | Exon+reg. EigenPhred | chr2:233701900 | rs182385495 | C/A | intron_variant | 0.0031 | 23 | 2.708758653 | 3.02E-02 |
| Bilirubin | *UGT1A9* | Exon+reg. EigenPhred | chr2:233755833 | . | G/C | non_coding_transcript_exon_variant | 0.0001 | 1 | 1.017775772 | 3.89E-01 |
| Bilirubin | *UGT1A9* | Exon+reg. EigenPhred | chr2:233712244 | rs184398589 | G/C | intron_variant | 0.0032 | 24 | 0.341924912 | 2.75E-01 |
| Bilirubin | *UGT1A9* | Exon+reg. EigenPhred | chr2:233692549 | . | T/C | intron_variant | 0.0001 | 1 | 1.712811794 | 9.55E-01 |
| Bilirubin | *UGT1A9* | Exon+reg. EigenPhred | chr2:233696404 | . | G/C | intron_variant | 0.0001 | 1 | 0.036079777 | 4.12E-01 |
| Bilirubin | *UGT1A9* | Exon+reg. EigenPhred | chr2:233744818 | . | T/C | intron_variant | 0.0001 | 1 | 0.461440961 | 9.06E-01 |
| Bilirubin | *UGT1A9* | Exon+reg. EigenPhred | chr2:233697534 | . | T/C | intron_variant | 0.0001 | 1 | 3.497236973 | 5.18E-01 |
| Bilirubin | *UGT1A9* | Exon+reg. EigenPhred | chr2:233758329 | rs544542552 | G/C | intron_variant | 0.0003 | 2 | 8.728609042 | 6.73E-01 |
| Bilirubin | *UGT1A9* | Exon+reg. EigenPhred | chr2:233714142 | . | A/G | intron_variant | 0.0001 | 1 | 0.145684728 | 3.12E-01 |
| Bilirubin | *UGT1A9* | Exon+reg. EigenPhred | chr2:233674345 | rs45539636 | T/C | intron_variant | 0.005 | 37 | 3.264489243 | 8.19E-01 |
| Bilirubin | *UGT1A9* | Exon+reg. EigenPhred | chr2:233692843 | rs746217324 | A/G | intron_variant | 0.0001 | 1 | 18.53452976 | 7.41E-01 |
| Bilirubin | *UGT1A9* | Exon+reg. EigenPhred | chr2:233766045 | . | T/C | intron_variant | 0.0001 | 1 | 2.418740181 | 9.78E-01 |
| Bilirubin | *UGT1A9* | Exon+reg. EigenPhred | chr2:233674318 | . | G/C | intron_variant | 0.0001 | 1 | 2.37966525 | 1.33E-01 |
| Bilirubin | *UGT1A9* | Exon+reg. EigenPhred | chr2:233712569 | . | G/C | intron_variant | 0.0001 | 1 | 3.876673167 | 3.84E-01 |
| Bilirubin | *UGT1A9* | Exon+reg. EigenPhred | chr2:233692033 | . | G/C | 3_prime_UTR_variant | 0.0001 | 1 | 9.088317071 | 9.12E-01 |
| Bilirubin | *UGT1A9* | Exon+reg. EigenPhred | chr2:233761970 | . | T/C | intron_variant | 0.0001 | 1 | 4.666034783 | 2.47E-01 |
| Bilirubin | *UGT1A9* | Exon+reg. EigenPhred | chr2:233767178 | rs753056825 | T/C | intron_variant | 0.0001 | 1 | 9.34144523 | 5.07E-01 |
| Bilirubin | *UGT1A9* | Exon+reg. EigenPhred | chr2:233714359 | rs754648080 | G/T | intron_variant | 0.0001 | 1 | 3.249637593 | 7.64E-01 |
| Bilirubin | *UGT1A9* | Exon+reg. EigenPhred | chr2:233672481 | rs145038612 | T/G | missense_variant | 0.0008 | 6 | 5.73524263 | 4.34E-01 |

| | | | | | | | | | | |
|---|---|---|---|---|---|---|---|---|---|---|
| Bilirubin | *UGT1A9* | Exon+reg. EigenPhred | chr2:233767878 | rs747062491 | A/T | synonymous_variant | 0.0003 | 2 | 11.51358054 | 7.74E-01 |
| Bilirubin | *UGT1A9* | Exon+reg. EigenPhred | chr2:233747198 | rs375624902 | C/T | intron_variant | 0.0003 | 2 | 0.452924555 | 5.17E-01 |
| Bilirubin | *UGT1A9* | Exon+reg. EigenPhred | chr2:233691799 | . | C/A | 5_prime_UTR_variant | 0.0001 | 1 | 9.670307217 | 9.87E-02 |
| Bilirubin | *UGT1A9* | Exon+reg. EigenPhred | chr2:233692049 | rs17868327 | C/A | 3_prime_UTR_variant | 0.0193 | 144 | 2.58984841 | 8.90E-02 |
| Bilirubin | *UGT1A9* | Exon+reg. EigenPhred | chr2:233712639 | rs184142486 | C/T | intron_variant | 0.0001 | 1 | 5.809753582 | 9.57E-01 |
| Bilirubin | *UGT1A9* | Exon+reg. EigenPhred | chr2:233712384 | rs45489204 | G/A | intron_variant | 0.0008 | 6 | 4.244844655 | 7.25E-01 |
| Bilirubin | *UGT1A9* | Exon+reg. EigenPhred | chr2:233697582 | rs139693283 | T/C | intron_variant | 0.0062 | 46 | 0.445700361 | 6.15E-01 |
| Bilirubin | *UGT1A9* | Exon+reg. EigenPhred | chr2:233692633 | . | C/T | intron_variant | 0.0001 | 1 | 5.625854125 | 7.35E-01 |
| Bilirubin | *UGT1A9* | Exon+reg. EigenPhred | chr2:233704993 | . | G/A | intron_variant | 0.0001 | 1 | 0.227624044 | 8.59E-01 |
| Bilirubin | *UGT1A9* | Exon+reg. EigenPhred | chr2:233672116 | . | A/T | missense_variant | 0.0001 | 1 | 6.384036323 | 9.27E-01 |
| Bilirubin | *UGT1A9* | Exon+reg. EigenPhred | chr2:233736992 | rs193257830 | G/T | intron_variant | 0.0017 | 13 | 0.895637889 | 2.21E-01 |
| Bilirubin | *UGT1A9* | Exon+reg. EigenPhred | chr2:233692288 | rs45477996 | C/T | 3_prime_UTR_variant | 0.0001 | 1 | 4.280105467 | 9.57E-01 |
| Bilirubin | *UGT1A9* | Exon+reg. EigenPhred | chr2:233761273 | rs748388612 | T/A | intron_variant | 0.0004 | 3 | 5.48085795 | 1.43E-01 |
| Bilirubin | *UGT1A9* | Exon+reg. EigenPhred | chr2:233761544 | rs745620081 | A/G | intron_variant | 0.0003 | 2 | 0.178036957 | 6.87E-01 |
| Bilirubin | *UGT1A9* | Exon+reg. EigenPhred | chr2:233737049 | . | A/G | intron_variant | 0.0001 | 1 | 0.199560044 | 2.36E-01 |
| Bilirubin | *UGT1A9* | Exon+reg. EigenPhred | chr2:233755691 | . | G/C | non_coding_transcript_exon_variant | 0.0001 | 1 | 1.104021668 | 1.95E-01 |
| Bilirubin | *UGT1A9* | Exon+reg. EigenPhred | chr2:233709891 | . | C/G | intron_variant | 0.0001 | 1 | 0.238058616 | 9.45E-01 |
| Bilirubin | *UGT1A9* | Exon+reg. EigenPhred | chr2:233762210 | rs190920095 | G/A | intron_variant | 0.0003 | 2 | 1.851922634 | 7.69E-01 |
| Bilirubin | *UGT1A9* | Exon+reg. EigenPhred | chr2:233674100 | . | A/C | intron_variant | 0.0046 | 34 | 2.538240673 | 1.02E-02 |
| Bilirubin | *UGT1A9* | Exon+reg. EigenPhred | chr2:233696793 | rs781512293 | T/C | intron_variant | 0.0001 | 1 | 3.159183971 | 9.98E-01 |
| Bilirubin | *UGT1A9* | Exon+reg. EigenPhred | chr2:233762256 | rs529163977 | A/G | intron_variant | 0.0001 | 1 | 1.803835412 | 1.88E-01 |
| Bilirubin | *UGT1A9* | Exon+reg. EigenPhred | chr2:233755843 | . | A/G | non_coding_transcript_exon_variant | 0.0001 | 1 | 3.922022913 | 7.40E-01 |
| Bilirubin | *UGT1A9* | Exon+reg. EigenPhred | chr2:233705435 | . | C/T | intron_variant | 0.0003 | 2 | 1.592304211 | 1.80E-01 |
| Bilirubin | *UGT1A9* | Exon+reg. EigenPhred | chr2:233755706 | . | C/G | non_coding_transcript_exon_variant | 0.0001 | 1 | 0.84121986 | 4.80E-01 |
| Bilirubin | *UGT1A9* | Exon+reg. EigenPhred | chr2:233696056 | . | A/T | intron_variant | 0.0007 | 5 | 1.679640144 | 1.88E-01 |
| Bilirubin | *UGT1A9* | Exon+reg. EigenPhred | chr2:233736722 | rs149071654 | G/A | intron_variant | 0.0275 | 205 | 2.352616426 | 2.07E-16 |
| Bilirubin | *UGT1A9* | Exon+reg. EigenPhred | chr2:233697743 | . | A/T | intron_variant | 0.0001 | 1 | 0.045875597 | 2.48E-01 |
| Bilirubin | *UGT1A9* | Exon+reg. EigenPhred | chr2:233692401 | . | C/T | intron_variant | 0.0001 | 1 | 7.857943249 | 7.57E-01 |
| Bilirubin | *UGT1A9* | Exon+reg. EigenPhred | chr2:233705644 | . | G/T | intron_variant | 0.0001 | 1 | 0.410318602 | 9.23E-01 |
| Bilirubin | *UGT1A9* | Exon+reg. EigenPhred | chr2:233767876 | rs773195449 | C/A | missense_variant | 0.0001 | 1 | 13.59240664 | 9.57E-01 |
| Bilirubin | *UGT1A9* | Exon+reg. EigenPhred | chr2:233679551 | rs139988189 | T/G | intron_variant | 0.0008 | 6 | 1.00685226 | 6.82E-02 |
| Bilirubin | *UGT1A9* | Exon+reg. EigenPhred | chr2:233672357 | rs138301590 | T/G | synonymous_variant | 0.0007 | 5 | 9.979425144 | 9.94E-01 |
| Bilirubin | *UGT1A9* | Exon+reg. EigenPhred | chr2:233692518 | rs745597997 | G/A | intron_variant | 0.0001 | 1 | 1.27470608 | 6.72E-01 |
| Bilirubin | *UGT1A9* | Exon+reg. EigenPhred | chr2:233676702 | rs748281452 | C/T | intron_variant | 0.0004 | 3 | 4.368413667 | 8.39E-01 |
| Bilirubin | *UGT1A9* | Exon+reg. EigenPhred | chr2:233696492 | rs181370414 | G/A | intron_variant | 0.0001 | 1 | 1.783249105 | 9.39E-01 |
| Bilirubin | *UGT1A9* | Exon+reg. EigenPhred | chr2:233761035 | rs57307513 | T/C | missense_variant | 0.0009 | 7 | 0.014589659 | 8.76E-01 |
| Bilirubin | *UGT1A9* | Exon+reg. EigenPhred | chr2:233766284 | . | C/T | intron_variant | 0.0001 | 1 | 0.091519444 | 6.07E-01 |
| Bilirubin | *UGT1A9* | Exon+reg. EigenPhred | chr2:233691800 | . | T/G | 5_prime_UTR_variant | 0.0001 | 1 | 9.385667397 | 9.87E-02 |
| Bilirubin | *UGT1A9* | Exon+reg. EigenPhred | chr2:233711092 | rs138181984 | G/T | intron_variant | 0.0003 | 2 | 2.24773047 | 7.37E-01 |
| Bilirubin | *UGT1A9* | Exon+reg. EigenPhred | chr2:233766390 | rs11679312 | C/T | intron_variant | 0.0263 | 196 | 0.856509811 | 5.14E-15 |
| Bilirubin | *UGT1A9* | Exon+reg. EigenPhred | chr2:233674397 | . | A/G | intron_variant | 0.0001 | 1 | 1.691126899 | 3.99E-01 |
| Bilirubin | *UGT1A9* | Exon+reg. EigenPhred | chr2:233772279 | rs202172337 | T/C | missense_variant | 0.0001 | 1 | 4.478758826 | 1.97E-01 |
| Bilirubin | *UGT1A9* | Exon+reg. EigenPhred | chr2:233697004 | rs761877638 | C/T | intron_variant | 0.0001 | 1 | 1.72809805 | 7.51E-01 |
| Bilirubin | *UGT1A9* | Exon+reg. EigenPhred | chr2:233711250 | . | T/C | intron_variant | 0.0001 | 1 | 1.356565804 | 6.40E-01 |
| Bilirubin | *UGT1A9* | Exon+reg. EigenPhred | chr2:233695287 | rs775200577 | A/G | intron_variant | 0.0005 | 4 | 0.867831605 | 2.15E-02 |
| Bilirubin | *UGT1A9* | Exon+reg. EigenPhred | chr2:233772306 | rs200370335 | G/T | missense_variant | 0.0001 | 1 | 14.66557688 | 7.41E-01 |
| Bilirubin | *UGT1A9* | Exon+reg. EigenPhred | chr2:233672736 | rs376424202 | A/G | missense_variant | 0.0001 | 1 | 13.78350234 | 6.12E-01 |
| Bilirubin | *UGT1A9* | Exon+reg. EigenPhred | chr2:233762199 | rs532806601 | A/G | intron_variant | 0.0001 | 1 | 0.742771004 | 6.65E-01 |
| Bilirubin | *UGT1A9* | Exon+reg. EigenPhred | chr2:233744657 | . | C/T | intron_variant | 0.0038 | 28 | 0.488161751 | 7.27E-02 |
| Bilirubin | *UGT1A9* | Exon+reg. EigenPhred | chr2:233697925 | . | G/A | intron_variant | 0.0001 | 1 | 0.710702763 | 8.13E-01 |
| Bilirubin | *UGT1A9* | Exon+reg. EigenPhred | chr2:233712285 | . | C/A | intron_variant | 0.0003 | 2 | 0.067006665 | 6.33E-01 |
| Bilirubin | *UGT1A9* | Exon+reg. EigenPhred | chr2:233672110 | rs769146316 | T/G | missense_variant | 0.0004 | 3 | 0.00181496 | 7.49E-02 |
| Bilirubin | *UGT1A9* | Exon+reg. EigenPhred | chr2:233675110 | . | T/A | intron_variant | 0.0001 | 1 | 0.558392454 | 2.70E-01 |
| Bilirubin | *UGT1A9* | Exon+reg. EigenPhred | chr2:233747267 | . | C/T | non_coding_transcript_exon_variant | 0.0003 | 2 | 1.36908278 | 6.80E-01 |
| Bilirubin | *UGT1A9* | Exon+reg. EigenPhred | chr2:233714452 | . | G/A | intron_variant | 0.0001 | 1 | 0.807470857 | 1.98E-01 |
| Bilirubin | *UGT1A9* | Exon+reg. EigenPhred | chr2:233762146 | rs541150164 | C/G | intron_variant | 0.0003 | 2 | 1.451470059 | 4.53E-01 |
| Bilirubin | *UGT1A9* | Exon+reg. EigenPhred | chr2:233691755 | rs144640470 | T/A | 5_prime_UTR_variant | 0.0073 | 54 | 10.59754905 | 1.56E-02 |
| Bilirubin | *UGT1A9* | Exon+reg. EigenPhred | chr2:233712538 | rs752556206 | G/A | intron_variant | 0.0001 | 1 | 2.623538358 | 7.09E-01 |
| Bilirubin | *UGT1A9* | Exon+reg. EigenPhred | chr2:233709856 | . | G/A | intron_variant | 0.0001 | 1 | 2.816186092 | 3.20E-01 |
| Bilirubin | *UGT1A9* | Exon+reg. EigenPhred | chr2:233758422 | . | T/G | intron_variant | 0.0001 | 1 | 4.781110994 | 2.32E-01 |
| Bilirubin | *UGT1A9* | Exon+reg. EigenPhred | chr2:233692141 | rs762078643 | C/A | 3_prime_UTR_variant | 0.0001 | 1 | 4.326411052 | 8.47E-01 |
| Bilirubin | *UGT1A9* | Exon+reg. EigenPhred | chr2:233705440 | rs72986482 | G/A | intron_variant | 0.033 | 246 | 0.030205814 | 4.79E-10 |
| Bilirubin | *UGT1A9* | Exon+reg. EigenPhred | chr2:233772599 | . | C/A | 3_prime_UTR_variant | 0.0001 | 1 | 7.381604525 | 2.02E-01 |
| Bilirubin | *UGT1A9* | Exon+reg. EigenPhred | chr2:233673975 | . | G/C | intron_variant | 0.0003 | 2 | 6.208398168 | 2.80E-01 |
| Bilirubin | *UGT1A9* | Exon+reg. EigenPhred | chr2:233711384 | rs547291485 | G/C | intron_variant | 0.0121 | 90 | 3.089475697 | 1.38E-01 |
| Bilirubin | *UGT1A9* | Exon+reg. EigenPhred | chr2:233747227 | rs138794928 | C/T | non_coding_transcript_exon_variant | 0.0109 | 81 | 1.970680035 | 9.70E-04 |
| Bilirubin | *UGT1A9* | Exon+reg. EigenPhred | chr2:233672699 | . | G/A | synonymous_variant | 0.0001 | 1 | 2.942288344 | 2.95E-01 |
| Bilirubin | *UGT1A9* | Exon+reg. EigenPhred | chr2:233692035 | rs555086706 | G/T | 3_prime_UTR_variant | 0.0004 | 3 | 6.451396534 | 3.39E-01 |
| Bilirubin | *UGT1A9* | Exon+reg. EigenPhred | chr2:233692652 | . | A/G | intron_variant | 0.0001 | 1 | 6.996057075 | 5.87E-02 |
| Bilirubin | *UGT1A9* | Exon+reg. EigenPhred | chr2:233772395 | rs751579554 | G/A | missense_variant | 0.0001 | 1 | 12.08550072 | 2.99E-01 |
| Bilirubin | *UGT1A9* | Exon+reg. EigenPhred | chr2:233674401 | . | T/C | intron_variant | 0.0001 | 1 | 0.249060812 | 6.63E-01 |
| Bilirubin | *UGT1A9* | Exon+reg. EigenPhred | chr2:233702891 | rs768271776 | C/T | intron_variant | 0.0001 | 1 | 1.800732258 | 5.15E-02 |
| Bilirubin | *UGT1A9* | Exon+reg. EigenPhred | chr2:233705564 | rs529349858 | C/A | intron_variant | 0.0016 | 12 | 0.033147236 | 7.58E-01 |
| Bilirubin | *UGT1A9* | Exon+reg. EigenPhred | chr2:233697120 | . | C/T | intron_variant | 0.0005 | 4 | 0.540577624 | 6.42E-01 |
| Bilirubin | *UGT1A9* | Exon+reg. EigenPhred | chr2:233709621 | . | T/A | intron_variant | 0.0001 | 1 | 3.127451785 | 9.92E-01 |
| Bilirubin | *UGT1A9* | Exon+reg. EigenPhred | chr2:233680695 | . | G/A | intron_variant | 0.0001 | 1 | 1.727837151 | 5.92E-01 |
| Bilirubin | *UGT1A9* | Exon+reg. EigenPhred | chr2:233697134 | rs537171274 | A/G | intron_variant | 0.0004 | 3 | 1.681954924 | 6.24E-01 |
| Bilirubin | *UGT1A9* | Exon+reg. EigenPhred | chr2:233674141 | rs28969703 | G/A | intron_variant | 0.0001 | 1 | 2.586477114 | 1.95E-01 |
| Bilirubin | *UGT1A9* | Exon+reg. EigenPhred | chr2:233692189 | . | T/C | 3_prime_UTR_variant | 0.0001 | 1 | 3.315600724 | 3.25E-01 |
| Bilirubin | *UGT1A9* | Exon+reg. EigenPhred | chr2:233772466 | . | C/A | synonymous_variant | 0.0001 | 1 | 12.10082516 | 9.07E-01 |
| Bilirubin | *UGT1A9* | Exon+reg. EigenPhred | chr2:233692445 | . | G/A | intron_variant | 0.0001 | 1 | 1.76897919 | 8.64E-01 |

| | | | | | | | | | | |
|---|---|---|---|---|---|---|---|---|---|---|
| Bilirubin | *UGT1A9* | Exon+reg. EigenPhred | chr2:233747259 | . | G/C | non_coding_transcript_exon_variant | 0.0003 | 2 | 2.957924105 | 6.80E-01 |
| Bilirubin | *UGT1A9* | Exon+reg. EigenPhred | chr2:233697068 | . | G/T | intron_variant | 0.0001 | 1 | 2.532686394 | 6.43E-01 |
| Bilirubin | *UGT1A9* | Exon+reg. EigenPhred | chr2:233747015 | . | G/T | intron_variant | 0.0001 | 1 | 0.23315682 | 8.41E-01 |
| Bilirubin | *UGT1A9* | Exon+reg. EigenPhred | chr2:233695282 | . | T/C | intron_variant | 0.0004 | 3 | 0.093819375 | 7.23E-03 |
| Bilirubin | *UGT1A9* | Exon+reg. EigenPhred | chr2:233711084 | . | G/A | intron_variant | 0.0001 | 1 | 0.389989781 | 8.86E-01 |
| Bilirubin | *UGT1A9* | Exon+reg. EigenPhred | chr2:233704913 | rs575509473 | C/T | intron_variant | 0.0003 | 2 | 2.773655505 | 4.70E-01 |
| Bilirubin | *UGT1A9* | Exon+reg. EigenPhred | chr2:233744651 | . | A/T | intron_variant | 0.0038 | 28 | 0.158557562 | 7.27E-02 |
| Bilirubin | *UGT1A9* | Exon+reg. EigenPhred | chr2:233695188 | . | C/T | intron_variant | 0.0004 | 3 | 0.986799327 | 5.30E-01 |
| Bilirubin | *UGT1A9* | Exon+reg. EigenPhred | chr2:233709314 | . | G/A | intron_variant | 0.0007 | 5 | 2.762461086 | 4.42E-01 |
| Bilirubin | *UGT1A9* | Exon+reg. EigenPhred | chr2:233695746 | . | A/G | intron_variant | 0.0001 | 1 | 0.93196606 | 8.69E-01 |
| Bilirubin | *UGT1A9* | Exon+reg. EigenPhred | chr2:233691866 | rs45464992 | A/G | 3_prime_UTR_variant | 0.0077 | 57 | 10.44894916 | 3.56E-03 |
| Bilirubin | *UGT1A9* | Exon+reg. EigenPhred | chr2:233755319 | rs776967353 | C/T | non_coding_transcript_exon_variant | 0.0001 | 1 | 6.394121005 | 7.36E-02 |
| Bilirubin | *UGT1A9* | Exon+reg. EigenPhred | chr2:233767069 | rs778321344 | T/C | synonymous_variant | 0.0001 | 1 | 17.20011978 | 6.07E-01 |
| Bilirubin | *UGT1A9* | Exon+reg. EigenPhred | chr2:233744714 | . | G/C | intron_variant | 0.0001 | 1 | 3.13845144 | 1.02E-01 |
| Bilirubin | *UGT1A9* | Exon+reg. EigenPhred | chr2:233691730 | . | A/T | 5_prime_UTR_variant | 0.0001 | 1 | 4.377461518 | 2.40E-01 |
| Bilirubin | *UGT1A9* | Exon+reg. EigenPhred | chr2:233675454 | . | G/C | intron_variant | 0.0003 | 2 | 0.072121858 | 5.17E-01 |
| Bilirubin | *UGT1A9* | Exon+reg. EigenPhred | chr2:233705549 | . | T/C | intron_variant | 0.0004 | 3 | 2.768628472 | 5.08E-01 |
| Bilirubin | *UGT1A9* | Exon+reg. EigenPhred | chr2:233702234 | . | A/G | intron_variant | 0.0001 | 1 | 1.691584614 | 4.56E-01 |
| Bilirubin | *UGT1A9* | Exon+reg. EigenPhred | chr2:233767154 | . | A/C | missense_variant | 0.0003 | 2 | 11.33117795 | 4.85E-01 |
| Bilirubin | *UGT1A9* | Exon+reg. EigenPhred | chr2:233755438 | rs192713031 | C/G | non_coding_transcript_exon_variant | 0.0003 | 2 | 5.935733997 | 7.37E-01 |
| Bilirubin | *UGT1A9* | Exon+reg. EigenPhred | chr2:233692193 | . | G/C | 3_prime_UTR_variant | 0.0001 | 1 | 9.091050072 | 9.31E-01 |
| Bilirubin | *UGT1A9* | Exon+reg. EigenPhred | chr2:233702910 | rs78155632 | T/A | intron_variant | 0.0079 | 59 | 0.404147222 | 1.55E-01 |
| Bilirubin | *UGT1A9* | Exon+reg. EigenPhred | chr2:233705497 | rs746321750 | C/T | intron_variant | 0.0004 | 3 | 1.191408434 | 7.68E-02 |
| Bilirubin | *UGT1A9* | Exon+reg. EigenPhred | chr2:233711404 | rs45498502 | C/T | intron_variant | 0.007 | 52 | 0.935700262 | 2.24E-01 |
| Bilirubin | *UGT1A9* | Exon+reg. EigenPhred | chr2:233760827 | rs148755655 | A/G | synonymous_variant | 0.0039 | 29 | 10.85555097 | 1.76E-01 |
| Bilirubin | *UGT1A9* | Exon+reg. EigenPhred | chr2:233674073 | rs536314636 | C/T | intron_variant | 0.0001 | 1 | 2.280983072 | 5.53E-01 |
| Bilirubin | *UGT1A9* | Exon+reg. EigenPhred | chr2:233744493 | . | T/G | intron_variant | 0.0001 | 1 | 1.005917431 | 6.72E-01 |
| Bilirubin | *UGT1A9* | Exon+reg. EigenPhred | chr2:233680696 | . | T/G | intron_variant | 0.0001 | 1 | 1.504935737 | 5.92E-01 |
| Bilirubin | *UGT1A9* | Exon+reg. EigenPhred | chr2:233747228 | . | C/G | non_coding_transcript_exon_variant | 0.0001 | 1 | 3.191576997 | 3.99E-01 |
| Bilirubin | *UGT1A9* | Exon+reg. EigenPhred | chr2:233692258 | rs754957875 | T/C | 3_prime_UTR_variant | 0.0003 | 2 | 3.283610752 | 3.37E-01 |
| Bilirubin | *UGT1A9* | Exon+reg. EigenPhred | chr2:233702350 | rs764503296 | T/A | intron_variant | 0.0005 | 4 | 2.223922055 | 5.64E-01 |
| Bilirubin | *UGT1A9* | Exon+reg. EigenPhred | chr2:233712596 | . | T/C | intron_variant | 0.0001 | 1 | 0.630008232 | 4.49E-01 |
| Bilirubin | *UGT1A9* | Exon+reg. EigenPhred | chr2:233767881 | . | G/A | synonymous_variant | 0.0001 | 1 | 5.675235789 | 7.04E-01 |
| Bilirubin | *UGT1A9* | Exon+reg. EigenPhred | chr2:233744499 | rs751170063 | T/C | intron_variant | 0.0003 | 2 | 0.601768182 | 3.91E-01 |
| Bilirubin | *UGT1A9* | Exon+reg. EigenPhred | chr2:233766036 | . | G/A | intron_variant | 0.0001 | 1 | 0.99134992 | 7.43E-02 |
| Bilirubin | *UGT1A9* | Exon+reg. EigenPhred | chr2:233768310 | . | C/T | missense_variant | 0.0003 | 2 | 14.62886565 | 8.83E-04 |
| Bilirubin | *UGT1A9* | Exon+reg. EigenPhred | chr2:233761943 | rs542276464 | G/A | intron_variant | 0.0028 | 21 | 7.591164732 | 1.18E-02 |
| Bilirubin | *UGT1A9* | Exon+reg. EigenPhred | chr2:233768291 | rs143573365 | G/A | missense_variant | 0.0003 | 2 | 10.6266056 | 7.30E-01 |
| Bilirubin | *UGT1A9* | Exon+reg. EigenPhred | chr2:233679323 | rs766783928 | C/T | intron_variant | 0.0001 | 1 | 0.097347383 | 1.52E-01 |
| Bilirubin | *UGT1A9* | Exon+reg. EigenPhred | chr2:233672201 | . | G/T | missense_variant | 0.0001 | 1 | 0.10766947 | 7.52E-02 |
| Bilirubin | *UGT1A9* | Exon+reg. EigenPhred | chr2:233709336 | . | A/G | intron_variant | 0.0001 | 1 | 0.073004681 | 4.86E-01 |
| Bilirubin | *UGT1A9* | Exon+reg. EigenPhred | chr2:233691852 | rs572506516 | G/A | 3_prime_UTR_variant | 0.0001 | 1 | 2.644802427 | 6.68E-01 |
| Bilirubin | *UGT1A9* | Exon+reg. EigenPhred | chr2:233709447 | rs756561648 | T/C | intron_variant | 0.0001 | 1 | 0.242675117 | 4.60E-01 |
| Bilirubin | *UGT1A9* | Exon+reg. EigenPhred | chr2:233767948 | rs112628426 | G/A | intron_variant | 0.0001 | 1 | 8.328302636 | 5.78E-01 |
| Bilirubin | *UGT1A9* | Exon+reg. EigenPhred | chr2:233761914 | rs28900396 | T/C | intron_variant | 0.0009 | 7 | 4.549307578 | 8.80E-01 |
| Bilirubin | *UGT1A9* | Exon+reg. EigenPhred | chr2:233695327 | rs372651727 | C/T | intron_variant | 0.0003 | 2 | 0.08072743 | 5.37E-01 |
| Bilirubin | *UGT1A9* | Exon+reg. EigenPhred | chr2:233711149 | rs562452811 | C/T | intron_variant | 0.0009 | 7 | 2.066835316 | 2.48E-01 |
| Bilirubin | *UGT1A9* | Exon+reg. EigenPhred | chr2:233671942 | rs145084767 | G/A | missense_variant | 0.0011 | 8 | 0.005571343 | 9.08E-02 |
| Bilirubin | *UGT1A9* | Exon+reg. EigenPhred | chr2:233702768 | . | G/C | intron_variant | 0.0001 | 1 | 2.253461789 | 5.78E-01 |
| Bilirubin | *UGT1A9* | Exon+reg. EigenPhred | chr2:233760569 | rs146052898 | T/C | synonymous_variant | 0.0001 | 1 | 5.286876813 | 5.41E-01 |
| Bilirubin | *UGT1A9* | Exon+reg. EigenPhred | chr2:233697327 | . | T/C | intron_variant | 0.0001 | 1 | 6.067581957 | 9.28E-01 |
| Bilirubin | *UGT1A9* | Exon+reg. EigenPhred | chr2:233695366 | rs184636578 | C/T | intron_variant | 0.0004 | 3 | 0.427766658 | 3.30E-01 |
| Bilirubin | *UGT1A9* | Exon+reg. EigenPhred | chr2:233744768 | rs116261344 | G/A | intron_variant | 0.0275 | 205 | 7.482900114 | 2.07E-16 |
| Bilirubin | *UGT1A9* | Exon+reg. EigenPhred | chr2:233674143 | . | T/A | intron_variant | 0.0001 | 1 | 3.326502216 | 3.03E-01 |
| Bilirubin | *UGT1A9* | Exon+reg. EigenPhred | chr2:233757461 | rs181439880 | C/T | intron_variant | 0.0011 | 8 | 3.433130341 | 7.79E-01 |
| Bilirubin | *UGT1A9* | Exon+reg. EigenPhred | chr2:233766694 | . | C/G | intron_variant | 0.0001 | 1 | 3.432102162 | 3.04E-01 |
| Bilirubin | *UGT1A9* | Exon+reg. EigenPhred | chr2:233766402 | . | C/T | intron_variant | 0.0001 | 1 | 1.241084545 | 7.46E-01 |
| Bilirubin | *UGT1A9* | Exon+reg. EigenPhred | chr2:233692030 | . | C/T | 3_prime_UTR_variant | 0.0003 | 2 | 7.645495195 | 3.74E-01 |

**Supplementary Table 4**
**Burden test *P*-values for adiponectin levels in the *ADIPOQ* gene, conditioned on known adiponectin and diabetes-associated variants.**

| rsID | position (GRCh38) on chromosome 3 | previous association | conditioned burden *P*-value |
|---|---|---|---|
| rs16861329 | 186948673 | type 2 diabetes | 4.76E-08 |
| rs17366568 | 186852664 | adiponectin levels | 3.79E-08 |
| rs182052 | 186842993 | adiponectin levels | 4.79E-08 |
| rs822387 | 186838248 | adiponectin levels, with and without BMI adjustment | 2.56E-07 |
| rs864265 | 186836503 | adiponectin levels | 5.56E-08 |
| rs1648707 | 186833922 | adiponectin levels | 4.68E-08 |
| rs10937273 | 186831906 | adiponectin levels | 5.24E-08 |
| rs6810075 | 186830776 | adiponectin levels | 4.77E-08 |
| rs266717 | 186812695 | adiponectin levels | 4.46E-08 |
| rs266719 | 186783859 | adiponectin levels | 2.40E-07 |
| rs822354 | 186762417 | adiponectin levels | 6.57E-08 |
| rs74577862 | 186843903 | adiponectin levels | 1.2E-07 |
| rs201813484 | 186841095 | adiponectin levels | 3.1E-07 |

**Supplementary Table 5.**
**Fraction of SNVs with |iHS|>2 in *APOC3*, *UGT1A9*, *ADIPOQ* and *FAM189B* compared to all other genes.** For each gene its fraction of SNVs with |iHS|>2 is given in parenthesis and the percentile in the empirical distribution of these fractions for all genes using four different definitions of the genomic region representing the genes. We mainly considered the most inclusive definition (the bottommost), but included the others for comparison to assess robustness to this definition. For *FAM189B* the percentile is also given for the subset of genes with a similar gene length, defined as the number of SNVs with iHS values (rightmost column). A percentile of 80% means that 80% of values are less than or equal to the value.

| **Definition of burden testing condition** | ***APOC3*** **compared to all genes** | ***UGT1A9*** **compared to all genes** | ***ADIPOQ*** **compared to all genes** | ***FAM189B*** **compared to all genes** | ***FAM189B*** **compared only to genes with within +/- 10% of # SNPs in *FAM189B*** |
|---|---|---|---|---|---|
| Exons only | 41.0th percentile (0.00) | 41.0th percentile (0.00) | 41.0th percentile (0.00) | 98.3th percentile (0.67) | 97.4th percentile (0.67) |
| Exons extended by 50bp and regulatory elements | 28.1th percentile (0.00) | 28.1th percentile (0.00) | 28.1th percentile (0.00) | 97.4th percentile (0.42) | 95.7th percentile (0.42) |
| Region spanning all exons | 15.3th percentile (0.00) | 15.3th percentile (0.00) | 15.3th percentile (0.00) | 96.7th percentile (0.32) | 94.6th percentile (0.32) |
| Region spanning all exons extended by 50bp and regulatory elements | 27.5th percentile (0.00) | 27.5th percentile (0.00) | 27.5th percentile (0.00) | 95.6th percentile (0.33) | 93.9th percentile (0.33) |

**Supplementary Table 6**
**Number of genes with at least 2 SNVs for the different burden analysis conditions.**

| Analysis condition | Number of genes |
|---|---|
| GENCODE V25 (all protein-coding, not tested) | 18,997 |
| LOFTEE HC | 85 |
| LOFTEE LC | 1,727 |
| Exon severe | 7,660 |
| Exon CADD | 18,428 |
| Exon CADD median | 18,138 |
| Exon+50 CADD | 18,551 |
| Exon+Regulatory Eigen | 18,961 |
| Exon+Regulatory EigenPhred | 18,660 |
| Exon+Regulatory EigenPCPhred | 18,722 |
| Regulatory only EigenPhred | 17,607 |

**Supplementary Table 7**
**Broad functional categories are defined by grouping together several variant categories as defined by Ensembl VEP.**

| Attributed category | Ensembl VEP functional class |
|---|---|
| intergenic_variant | intergenic_variant |
| intron_variant | intron_variant |
| UTR | 3_prime_UTR_variant |
| | 5_prime_UTR_variant |
| Up/Down stream | upstream_gene_variant |
| | downstream_gene_variant |
| splice_variant | splice_donor_variant |
| | splice_acceptor_variant |
| | splice_region_variant |
| coding_variant | coding_sequence_variant |
| | incomplete_terminal_codon_variant |
| | initiator_codon_variant |
| | missense_variant |
| | start_lost |
| | stop_gained |
| | stop_lost |
| | stop_retained_variant |
| | synonymous_variant |
| non_coding_variant | nc_transcript_variant |
| | non_coding_transcript_exon_variant |
| | non_coding_exon_variant |
| | mature_miRNA_variant |
| regulatory_variant | regulatory_region_variant |
| | TF_binding_site_variant |

**Supplementary Figure 1**

**Sequencing depth distribution of 1,457 MANOLIS samples.** The box indicates quartiles and the bold line is the median. Whiskers extend to 1.5x the interquartile range. The mean is 22.5x and the median is 21.9x. Sequencing depths range from 14.7x to 40x.

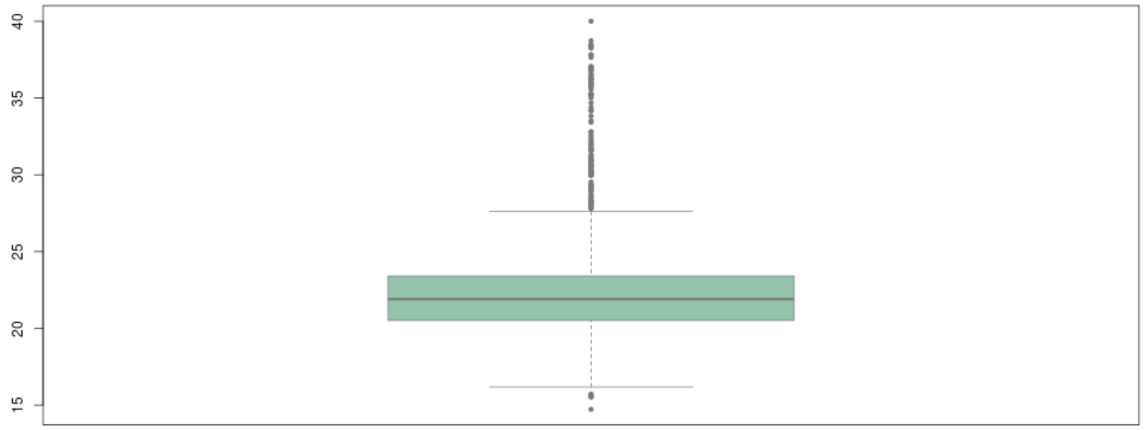

**Supplementary Figure 2**
**True positive rate for NA12878 based on Genome in a Bottle 0.2 at various sequencing depths for SNVs and INDELs.**

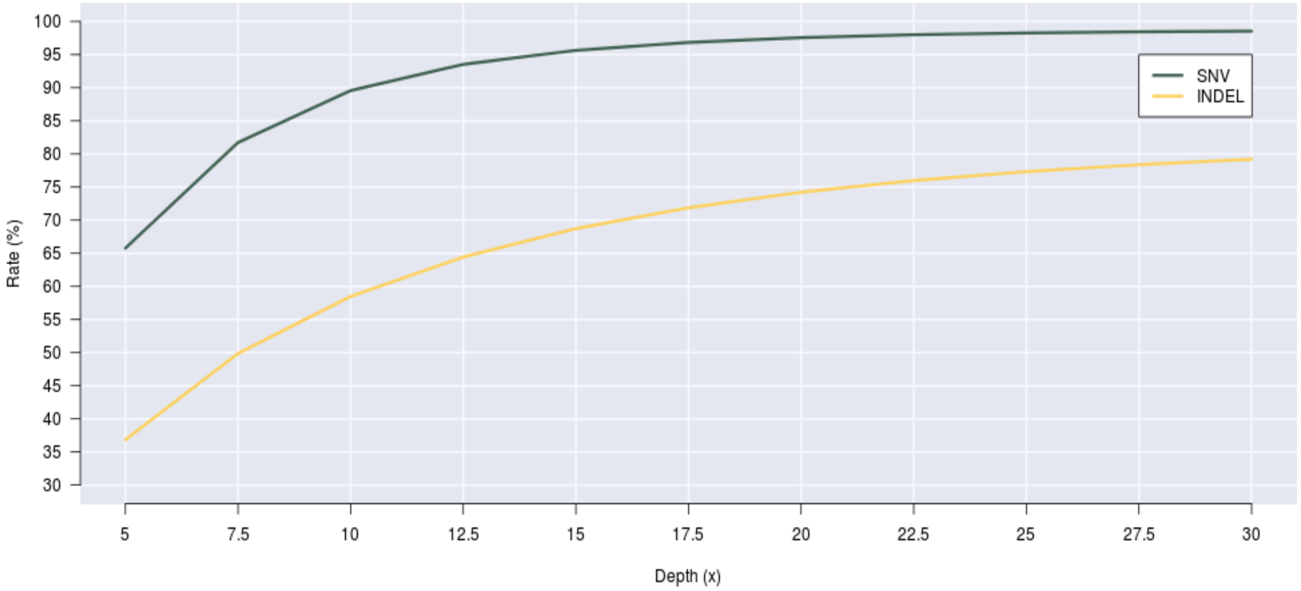

**Supplementary Figure 3**
**Breakdown of unique and shared SNVs between 30x and 22.5x depth whole genome sequencing callsets.**

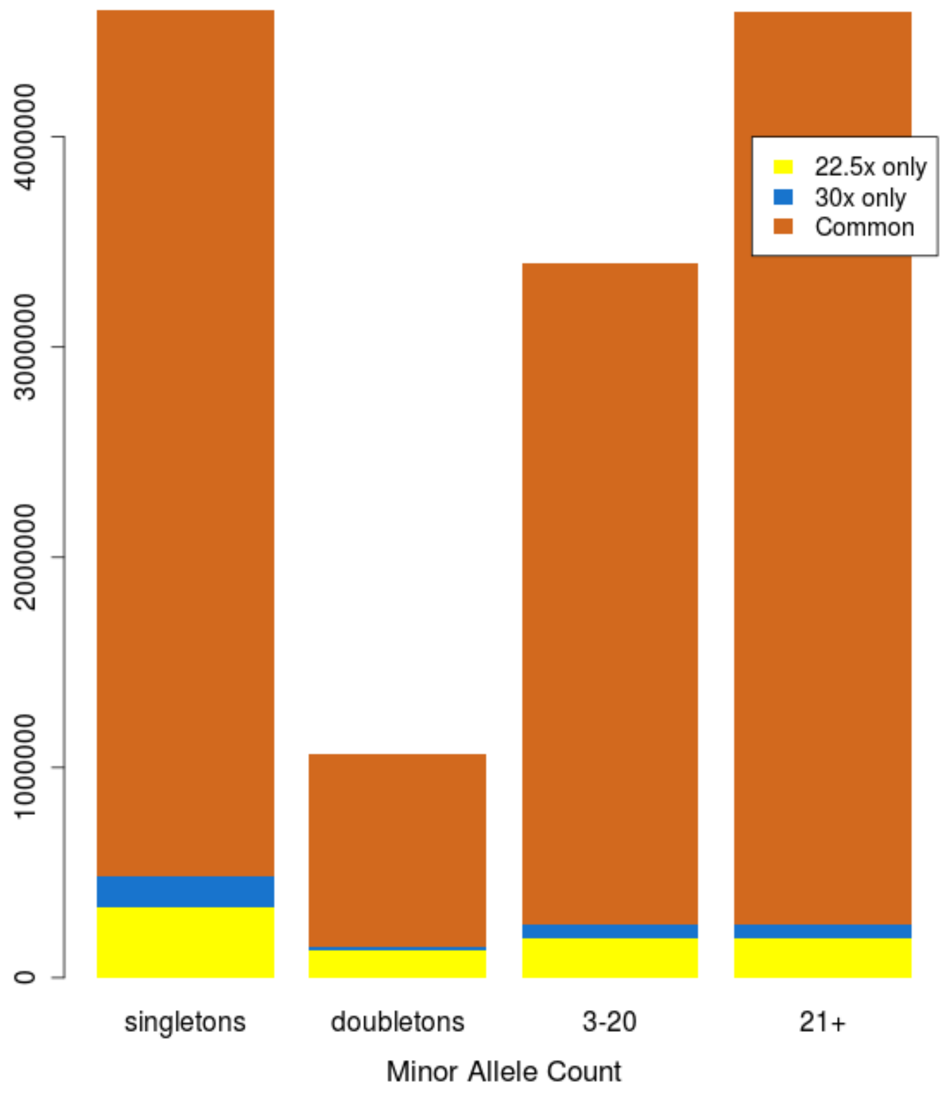

**Supplementary Figure 4**
**Variants predicted as loss-of-function with (a) high-confidence (HC) and (b) low-confidence (LC) by LOFTEE, genome-wide counts per Ensembl predicted consequence.**

**a.**

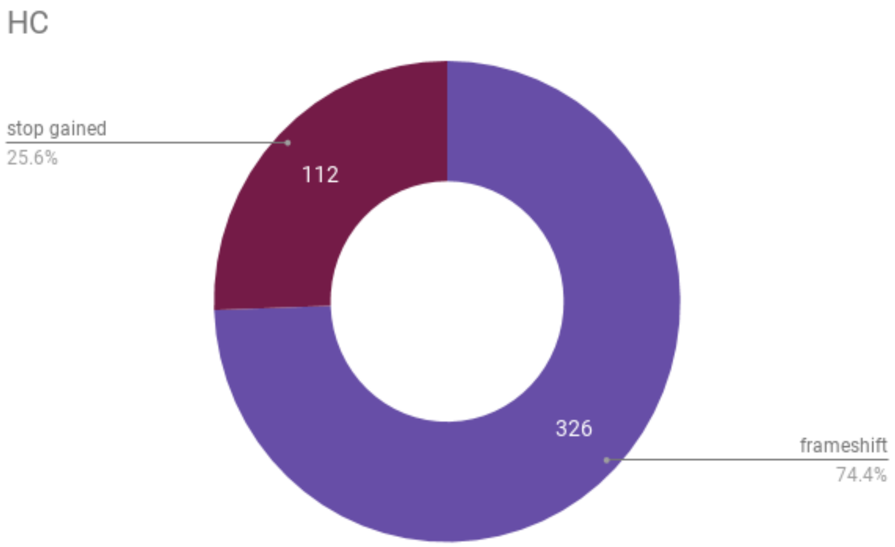


**b.**

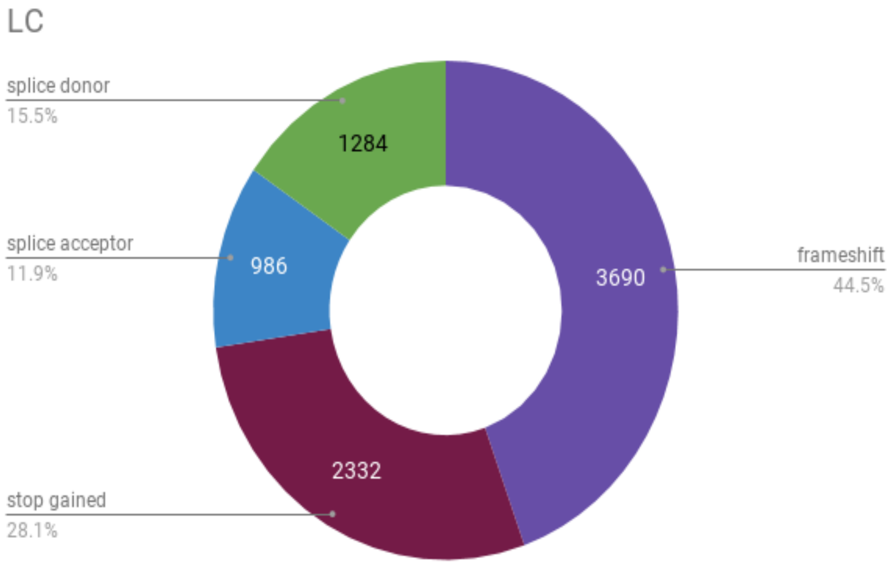

**Supplementary Figure 5**

**Distributions of (a) singleton and (b) doubleton counts in 1,000 draws of 100 MANOLIS samples.** The vertical orange line indicates the observed count in 100 TEENAGE samples downsampled to 22.5x.

**a.**

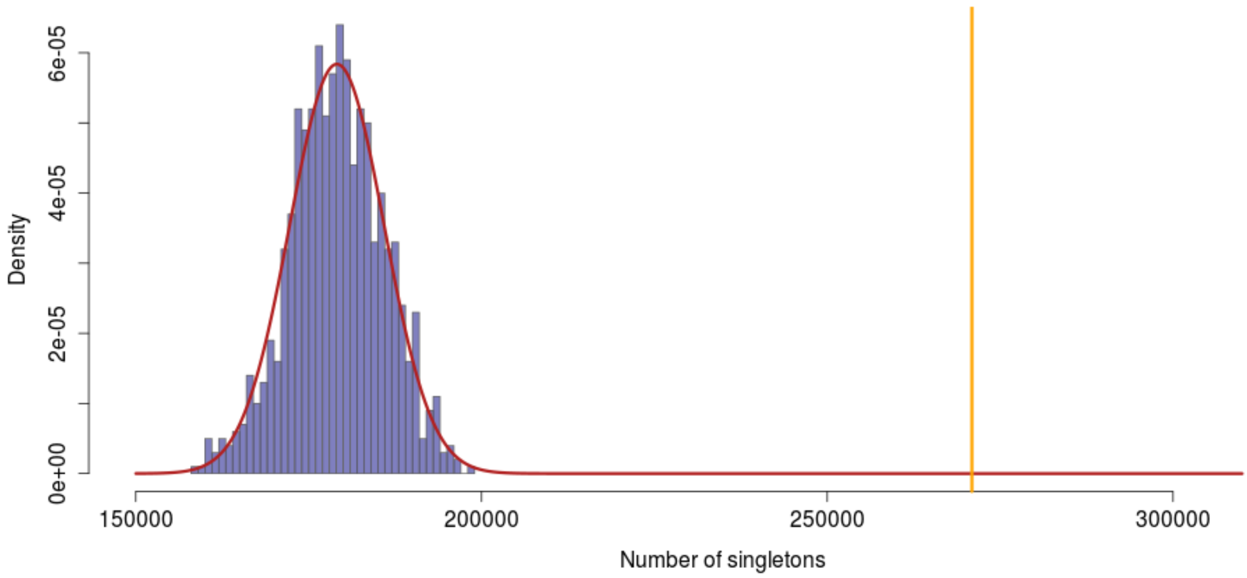


**b.**

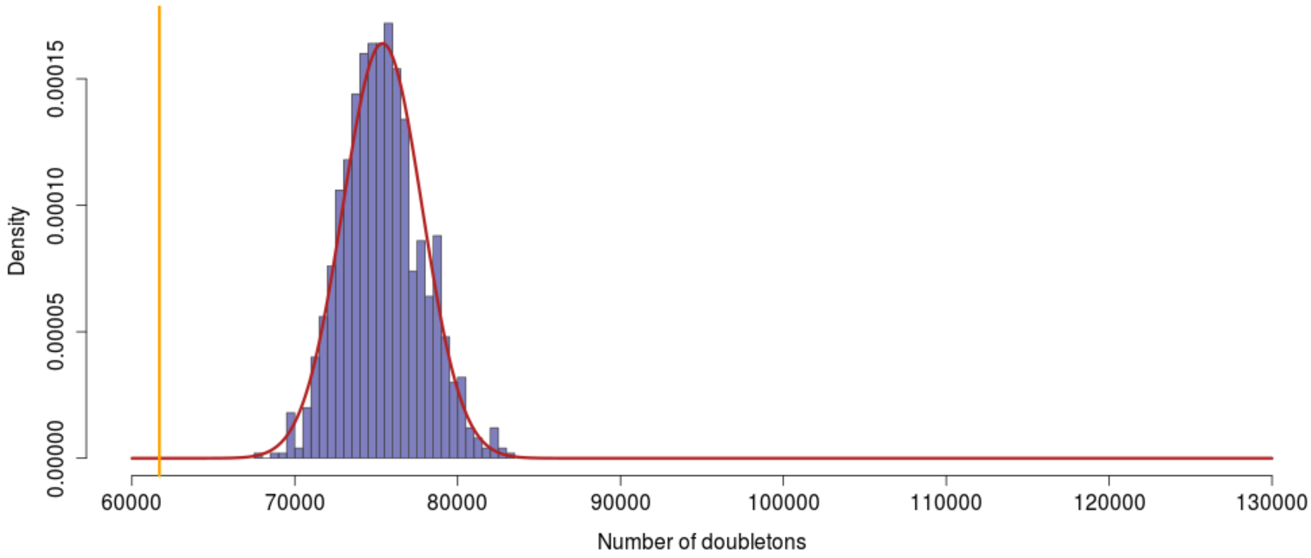

**Supplementary Figure 6**
**(a) Frequencies and (b) functional annotations of all novel variants in the MANOLIS whole genome sequence data.** Novelty is established by comparing variant location and alleles to Ensembl VEP annotation as well as gnomAD genomic variants lifted-over to build 38.

a.

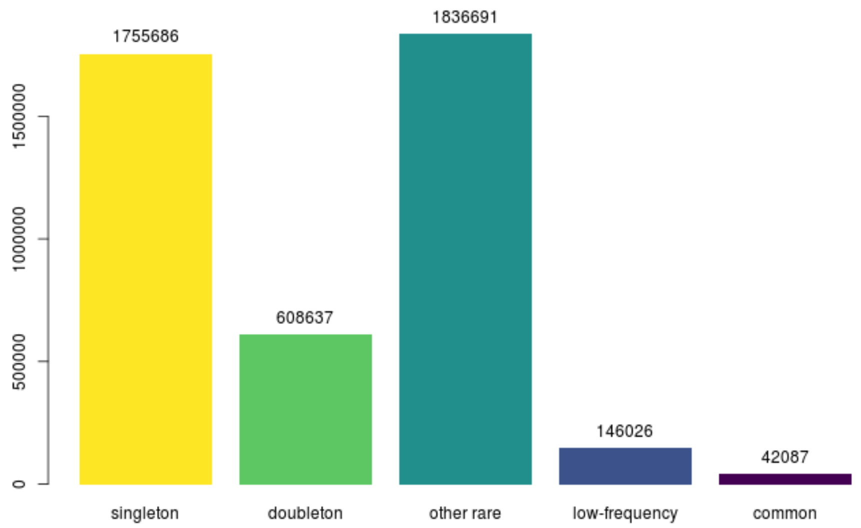


b.

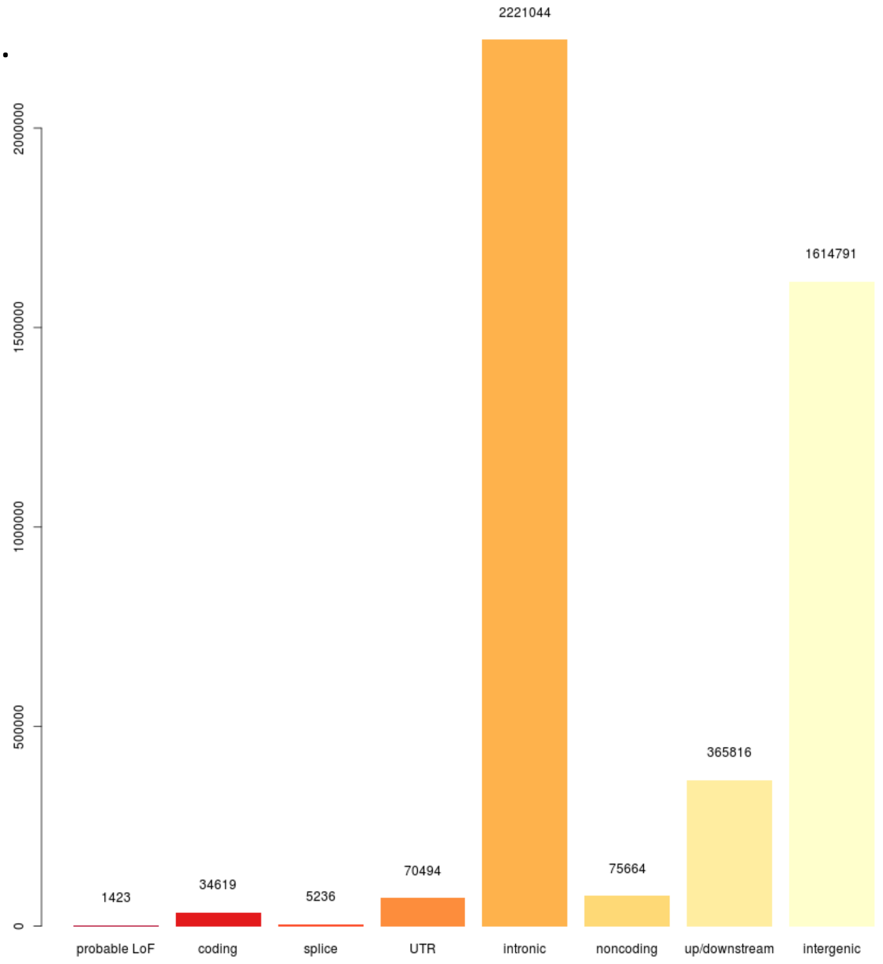

**Supplementary Figure 7**

**Correlogram of z-scores arising from all evaluated burden testing scenarios.** Quantities reported are Pearson's correlations of z-transformed *P*-values across all gene-trait pairs for all six tested traits. For each cell, *P*-values of all gene-trait pairs that were tested in both conditions are included. Clusters were generated using hierarchical clustering. The exon LoF HC scenario is not included in the correlogram due to the low number of genes containing more than one high-confidence loss-of-function variant (n=85).

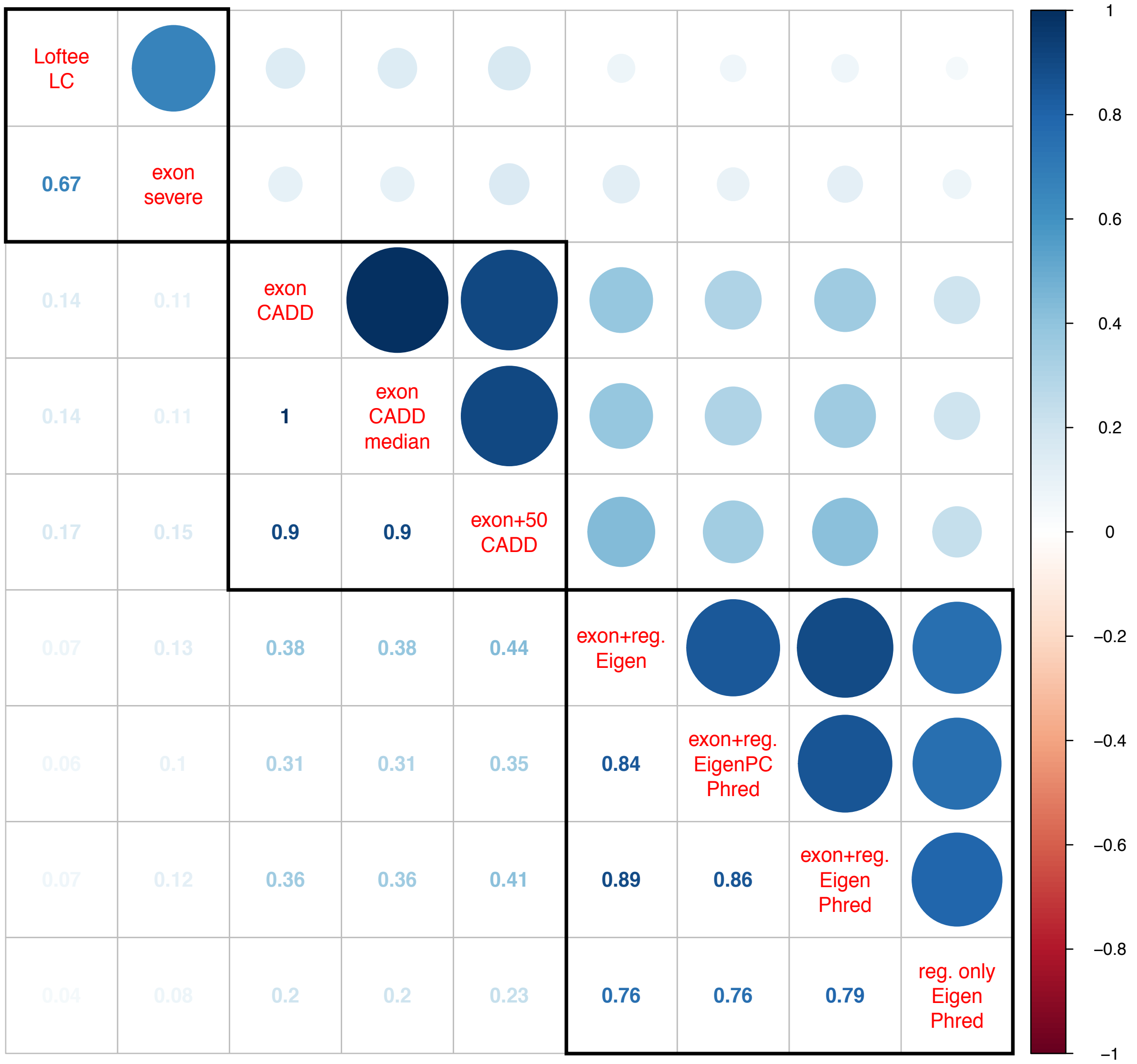

**Supplementary Figure 8**

**Evidence for association (*P*-values on the $-\log_{10}$ scale) for the 20 study-wide significant trait-gene pairs across all tested conditions.** Grey cells indicate that an insufficient number of variants passed the inclusion threshold. Dark orange denotes study-wide significance and turquoise green denotes suggestive association.

| | Exon | | | | | Exon+regulatory | | | reg. only |
|---|---|---|---|---|---|---|---|---|---|
| | Loftee LC | Exon severe | Exon CADD | Exon CADD+50 | Exon CADD median | Exon+Reg EigenPC Phred | Exon+Reg Eigen Phred | Exon+reg Eigen | Regulatory only EigenPhred |
| adiponectin *ADIPOQ* | | 2.72 | 5.03 | 5.7 | 5.03 | 6.43 | 7.27 | 7.37 | 0.708 |
| HDL *APOC3* | 20.3 | 19.7 | 14.3 | 19.5 | 14.3 | 2.54 | 5.65 | 8.09 | 5.76 |
| TG *FAM189B* | | 6.8 | 0.902 | 0.867 | 0.902 | 0.844 | 0.742 | 0.906 | 0.46 |
| Bilirubin *UGT1A10* | | | 0.238 | 1.3 | 0.238 | 6.66 | 7.24 | 7.69 | 7.47 |
| Bilirubin *UGT1A3* | | | 5.85 | 5.76 | 5.85 | 6.86 | 5.55 | 5.64 | 5.77 |
| Bilirubin *UGT1A6* | | | 2.37 | 7.08 | 2.37 | 7.77 | 7.03 | 7.69 | 7.0 |
| Bilirubin *UGT1A7* | | | 0.219 | 0.995 | 0.219 | 6.71 | 5.69 | 6.8 | 6.65 |
| Bilirubin *UGT1A8* | | | 0.629 | 1.02 | 0.629 | 6.6 | 7.27 | 7.25 | 7.47 |
| Bilirubin *UGT1A9* | | | 0.285 | 2.38 | 0.285 | 6.49 | 7.17 | 7.71 | 7.14 |
| Bilirubin *USP40* | 0.155 | 0.155 | 1.06 | 1.05 | 1.06 | 0.324 | 0.325 | 6.81 | 5.34 |

**Supplementary Figure 9**

**Comparison of kinship coefficients produced by KING and EMMAX to those produced by GEMMA[24]**. All kinship coefficients are calculated using the same dataset (MAF>5%, missingness<1%, LD-pruned). IBS coefficients (KING Related, EMMAX IBS and Plink) are both higher on average and less sensitive to increased relatedness than their Balding-Nichols counterparts (GEMMA, EMMAX BN and KING homogenous). Red lines represent per-dataset OLS regression slopes.

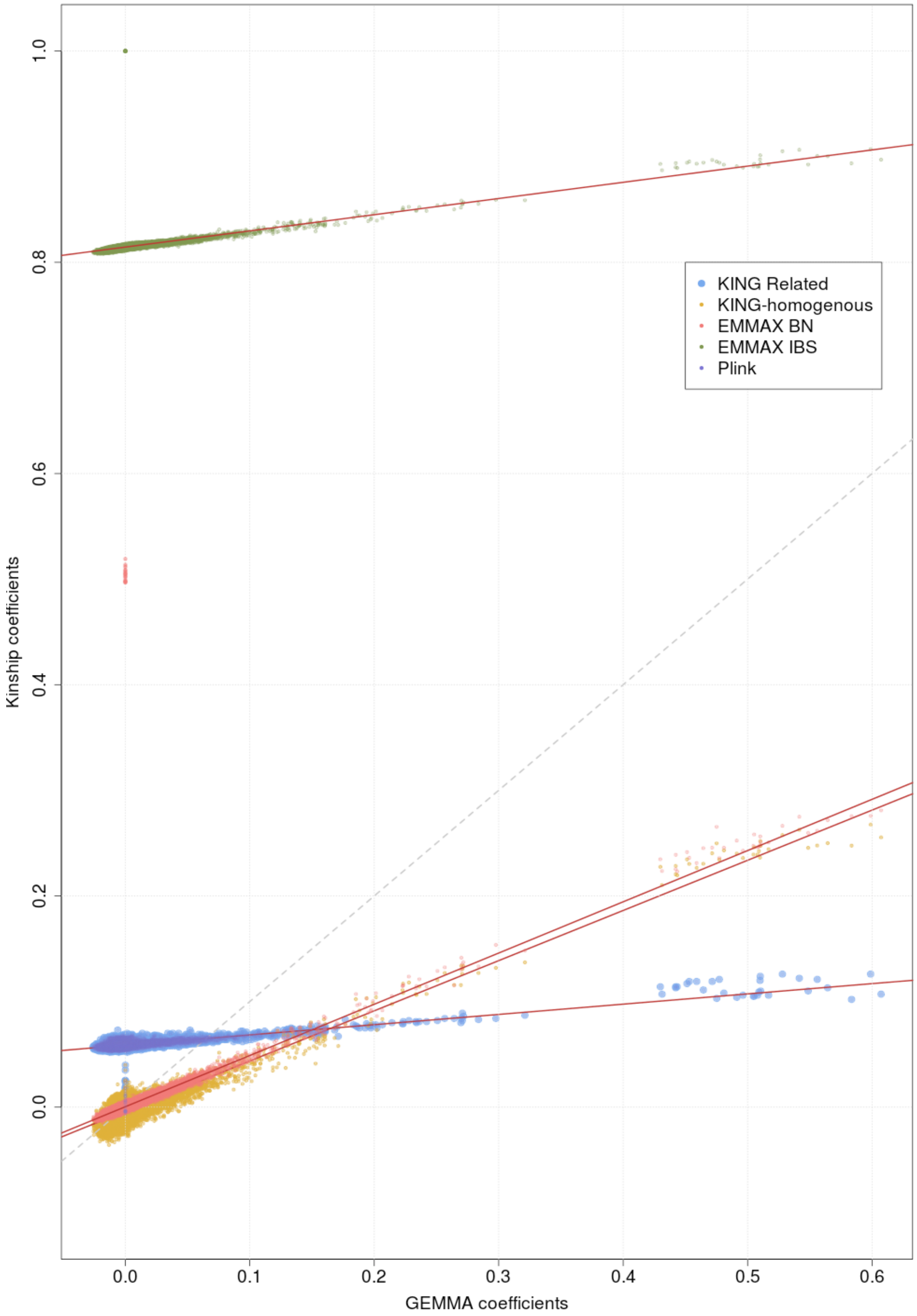
